# An apicomplexan-specific foundation model improves prediction of regulatory genomic signals in *Plasmodium falciparum*

**DOI:** 10.64898/2026.11.12.766050

**Authors:** Alan S. Brown, Jianyu Yang, Manuel Llinás, Shaun Mahony

**Affiliations:** Center for Eukaryotic Gene Regulation, Department of Biochemistry & Molecular Biology, The Pennsylvania State University, PA 16802, USA; Huck Center for Malaria Research, The Pennsylvania State University, PA 16802, USA; Department of Chemistry, The Pennsylvania State University, PA 16802, USA; Department of Computer Science and Engineering, The Pennsylvania State University, PA 16802, USA

## Abstract

Genomic foundation models learn transferable sequence representations through self-supervision, yet their practical value is contested. In data-rich mammalian genomes, task-specific models trained from scratch can match or exceed pretrained models on many regulatory prediction tasks. Whether self-supervised pretraining provides an advantage in small, data-sparse genomes has not been directly tested. We address this in the compact, 23Mbp, A/T-rich genome of the malaria parasite, *Plasmodium falciparum*. We trained a Genomic Pretrained Network (GPN) via masked-nucleotide reconstruction on 455 genome assemblies spanning 76 apicomplexan species. We then fine-tuned the model to predict a variety of epigenomic features in a held-out chromosome, including transcription factor binding sites, chromatin-accessibility, histone modifications, and 10 chromatin states. Pretraining followed by fine tuning improved prediction of most features, with most of the benefit concentrated on datasets with relatively little labeled data (fewer than 3,000 training examples). In contrast, a human-pretrained model transferred poorly, indicating that phylogenetically relevant pretraining, not foundation-model status alone, drives beneficial performance. Self-supervised pretraining on clade-specific assemblies is thus a practical route to data-efficient regulatory modeling in small genomes where training data is scarce.

## Introduction

Deep learning has become a central approach for predicting regulatory elements or gene expression directly from DNA sequence. Early convolutional neural networks (CNNs) demonstrated that short DNA windows contain rich information sufficient to predict diverse chromatin features and can predict the effects of non-coding variants (Kelley et al., 2018; J. Zhou & Troyanskaya, 2015). More recent architectures consider longer stretches of sequence when making each prediction, enabling higher performance on complex, long-range genomic tasks (Avsec et al., 2021, 2026). In parallel, genomic foundation models have emerged (Benegas et al., 2023, 2025; Brixi et al., 2026; Dalla-Torre et al., 2025; Nguyen et al., 2023). These models are first trained on large collections of unlabeled genome sequences (pretraining) and then adapted to specific prediction tasks using smaller labeled datasets (fine-tuning). By pretraining on diverse genome sequences, foundation models attempt to capture informative DNA features. Genomic foundation models have been fine-tuned to predict transcription factor (TF) binding, chromatin accessibility, and gene expression across a wide range of organisms including fungi, protozoa, mammals, and bacteria (Dalla-Torre et al., 2025; Ji et al., 2021; Nguyen et al., 2023; Z. Zhou et al., 2024).

Foundation model pretraining typically uses a self-supervised training task, where the training labels can be derived from the inputs themselves without the need for external labels. In masked nucleotide reconstruction, for example, several bases in each input sequence are hidden, and the model trains to predict the missing bases from the surrounding sequence context. Because the prediction target is already present in the original unmasked sequence, self-supervised pretraining procedures require only genome sequence information. DNABERT-2 was pretrained to reconstruct masked stretches of sequence from genomes of 135 species (Z. Zhou et al., 2024). The Genomic Pretrained Network (GPN), the model architecture we use here, was pretrained to reconstruct masked nucleotides from unlabeled plant genomes (Benegas et al., 2023). Evo 2 was pretrained to predict the next base in a sequence of nucleotides, using roughly 9 trillion base pairs from all domains of life (Brixi et al., 2026).

The premise of self-supervised pretraining is that, to predict hidden bases accurately, a model must learn the sequence patterns that recur within and across genomes, including protein-coding sequences, repetitive elements, and the regulatory grammar of TF binding motifs. If a model’s weights encode informative DNA features during pretraining, these weights could then be reused when the model is fine-tuned to specific downstream tasks, such as predicting if an input sequence is a TF binding site. Fine-tuning training processes reuse most of the core model architecture and weights that resulted from the pretraining step. The final model layers that were used to predict nucleotides in the self-supervised training task are replaced by classification layers, but most of the model architecture remains the same. The model is then trained using labeled data for the downstream task at hand. A major motivation driving the development of genomic foundation models is that if a pretrained model already encodes important sequence features, fine-tuned classifiers should perform more robustly or might need fewer labeled examples to learn the downstream task.

Despite the initial promise of foundation models, the practical value of pretraining has not been uniform across settings. Pretrained models have produced underwhelming results relative to randomly initialized models trained specifically for the downstream task (Z. Tang et al., 2025). Benchmarking efforts emphasize that the efficacy of foundation-model pretraining depends on the volume of labeled samples available, the training data construction settings, and the degree of domain shift between pretraining and fine-tuning data (Feng et al., 2025; Mupparapu et al., 2025; You et al., 2025). For human regulatory genomic prediction tasks, carefully trained task-specific models can match or exceed the performance of fine-tuned foundation models (Feng et al., 2025; Z. Tang et al., 2025; You et al., 2025). However, human regulatory genomic tasks are relatively data-rich, as mammalian-scale genomes offer numerous training samples and vast collections of functional genomics data exist from efforts like ENCODE (ENCODE Consortium, 2004). The application of foundation models to organisms with smaller genomes and smaller training datasets is under-explored. This motivates us to specifically assess pretraining strategies in cases where minimal labeled training data is available.

Single-celled eukaryotic apicomplexan parasites provide a biologically important setting in which to test genomic foundation model pretraining and fine-tuning methods. Several genera in this phylum cause diseases in humans or livestock, including *Toxoplasma*, *Plasmodium*, *Cryptosporidium*, *Babesia*, and *Theileria*. Apicomplexan genomes are also substantially different from the genomes used to train most existing foundation models. For example, the malaria parasite *Plasmodium falciparum* has an extremely high A/T content of 80.6% (Gardner et al., 2002). Its 23Mb genome also has several atypical associations between histone modifications and regulatory activities (Bartfai et al., 2010; Karmodiya et al., 2015; Salcedo-Amaya et al., 2009) and nucleosomes with reduced stability (Silberhorn et al., 2016; Watzlowik et al., 2021).

While gene expression patterns are highly dynamic across complex lifecycles in mosquito and human hosts, the *P. falciparum* genome encodes under 100 predicted sequence-specific TFs (Bischoff & Vaquero, 2010). A subset of these are the 30 identified members of the apicomplexan APETALA2 (ApiAP2) family (Balaji et al., 2005; Campbell et al., 2010; Singhal et al., 2024), which are attractive drug targets because they have no known homologs in humans (Jeninga et al., 2019; Russell et al., 2022; Singhal et al., 2024).

Because of the differences in gene regulatory architecture, we reasoned that existing foundation models trained on mammalian genomes, or those trained across a wide variety of eukaryotic genomes, would be unlikely to encapsulate regulatory features specific to apicomplexan genomes. We therefore trained a clade-specific foundation model using masked nucleotide reconstruction across collections of apicomplexan genome sequences. In contrast to the abundant data resources available for mammalian systems, regulatory prediction tasks in apicomplexans face acute limitations in the amount of labeled training data (Brown et al., 2026; Read et al., 2019). Despite the lack of available labeled training data, numerous genome sequence assemblies exist for this clade. The asymmetry between the abundance of available sequencing data and the scarcity of labeled datasets makes apicomplexan systems a natural domain where cross-species or cross-task pretraining could plausibly improve generalization as a basis for fine tuning in a data-sparse setting.

Realizing the potential benefits of pretrained models in low data settings depends on how the pretrained model is adapted to the downstream task. Available strategies differ mainly in how and to what degree the core model weights are updated (Ding et al., 2023; Lialin et al., 2024). The classic method of full fine-tuning updates all model weights. Full fine-tuning is computationally costly and can overwrite the representations that the pretraining produced (Kumar et al., 2022). Parameter-efficient methods instead hold the pretrained weights constant (“freezing” them) and update only a small set of added or selected parameters. Low-Rank Adaptation (LoRA) (Hu et al., 2021; Zhan et al., 2024; Z. Zhou et al., 2024), prompt tuning (Lester et al., 2021; X. L. Li & Liang, 2021; Nguyen et al., 2023), and Infused Adapter by Inhibiting and Amplifying Inner Activations (IA)^3^ (Dalla-Torre et al., 2025; Liu et al., 2022) are prominent examples of parameter-efficient fine-tuning techniques. Each of these methods attempts to prevent the model from forgetting the features it learned during pretraining. The simplest option is to freeze the model core entirely and train only lightweight classification layers on the fixed embedding space produced by the model core (Marin et al., 2023). Linear Probe then Fine Tune (LPFT) starts with freezing the model core then unfreezes the model core after half the training run to allow all model weights to update at a reduced learning rate (Kumar et al., 2022). The benefits of these strategies are expected to be largest precisely when labeled data are scarce, because that is when overwriting pretrained representations is the most costly. Establishing whether pretraining is beneficial and which adaptation strategy realizes that benefit therefore requires a domain that is data limited.

In this study, we assess whether self-supervised pretraining on apicomplexan genome sequences improves downstream prediction of regulatory genomic signals in *P. falciparum*. We evaluate self-supervised models directly to assess the degree to which they have captured regulatory features in the *P. falciparum* genome. We then compare the performance of models with and without pretraining across a range of regulatory genomics prediction tasks in *P. falciparum*, and we investigate settings under which foundation model-based pretraining is advantageous. Together, our experiments suggest that clade-specific foundation model pretraining can offer improved performance when predicting epigenomic signals in small genomes and sparse data settings.

## Results

### An apicomplexan-specific foundation model trained using all available genome sequences outperforms annotation-guided models

To build an apicomplexan-specific foundation model, we chose the CNN-based GPN architecture (Benegas et al., 2023) and a masked nucleotide reconstruction training task (**Fig. 1**) because CNNs have been shown to out-perform alternative architectures in related contexts (Z. Tang et al., 2025; Tay et al., 2022; K. K. Yang et al., 2024). We considered two distinct approaches to pretraining foundation models for use with downstream regulatory genomics prediction tasks. In the first approach, we pretrained on all 455 apicomplexan genome assemblies available in the National Center for Biotechnology Information (NCBI) database (**Supplemental Fig. 1**). We initially trained this model for 250,000 iterations (“Whole Genome 455A 250k steps”), but observing that training loss had not plateaued at that point, we continued training to a total of two million iterations (“Whole Genome 455A 2M steps”) (**Supplemental Fig. 2**). In the second approach, rationalizing that downstream tasks such as TF binding site prediction will benefit more from focusing pretraining on potential regulatory regions, we pretrained models using sequences around annotated transcription start sites (TSSs) (see **Methods**). RefSeq gene annotations are only available for 51 apicomplexan genome assemblies in the NCBI database (**Supplemental Fig. 1**). We trained two models that rely on these RefSeq annotations to demarcate genic regions: one model was trained using 1 kbp sequences upstream of TSSs and all exonic sequences (“Promoters+Exons 51A”); the second model was trained using 2 kbp sequence windows centered on annotated TSSs (“TSS Centered 51A”). Both annotation-based models were trained for 250,000 iterations.

**Figure 1.**
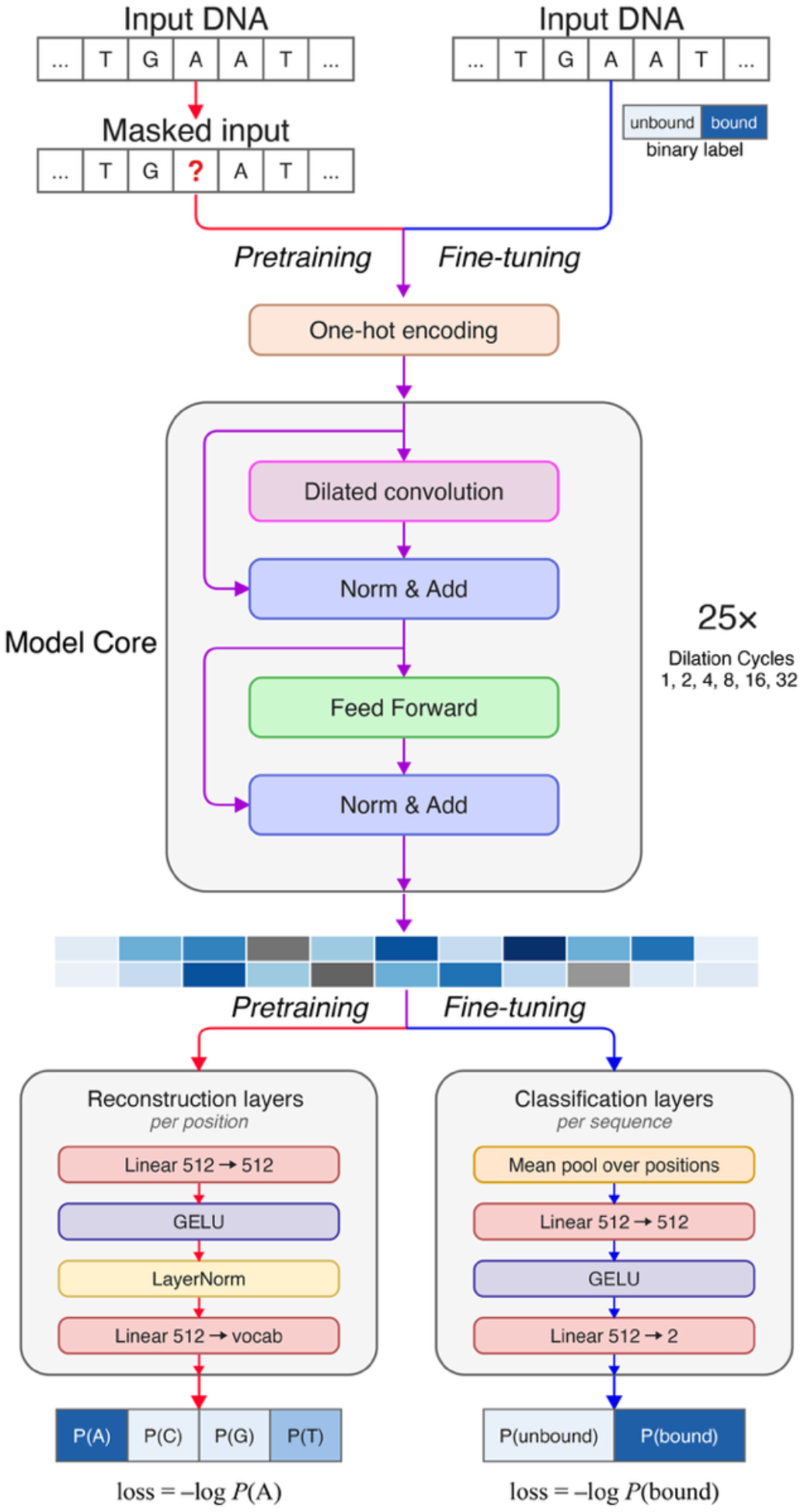
Model architecture for pretraining and classification tasks. A single convolutional model core (purple arrows) is shared by two different training pathways. Pretraining (red arrows): the model is trained to reconstruct masked nucleotides. Fine-tuning (blue arrows): *P. falciparum* sequences are given binary labels based on ChIP-seq, ATAC-seq, or chromatin state annotations. In both stages, sequences are one-hot-encoded and passed through 25 convolutional layers which output a 512-dimensional embedding at every position of the input sequence. That embedding is fed into either the reconstruction layers or the classification layers.

For testing, we withheld chromosome 14, the largest chromosome in the *P. falciparum* genome. To prevent models from merely memorizing sequences similar to the test chromosome, we aligned the *P. falciparum* chromosome 14 to all of the other training assemblies using Minimap2 (H. Li, 2018) and removed matching sequences from the training set. To assess how accurately each model can reconstruct the *P. falciparum* genome, we sequentially gave the model 512bp sequences from the *P. falciparum* genome with the center base masked, such that every base in the genome was masked and predicted. We saved the probability assigned to the correct base for every nucleotide in the *P. falciparum* genome.

The distribution of probabilities assigned by each model to the correct bases are shown in **Fig. 2A**. The “Whole Genome” models clearly outperform annotation-dependent models (i.e., “Promoters+Exons 51A” and “TSS Centered 51A”) in terms of accurately predicting masked bases, suggesting that models trained with a greater diversity of sequences can outperform models where training focuses on functional regions. The Whole Genome model trained for 2 million steps (i.e., “Whole Genome 455A 2M steps”) performs significantly better than the Whole Genome model (i.e., “Whole Genome 455A 250k steps”). The distributions of probabilities assigned to correct bases are very similar across training and test sets in all four models, suggesting minimal overfitting despite the small *P. falciparum* genome size.

**Figure 2.**
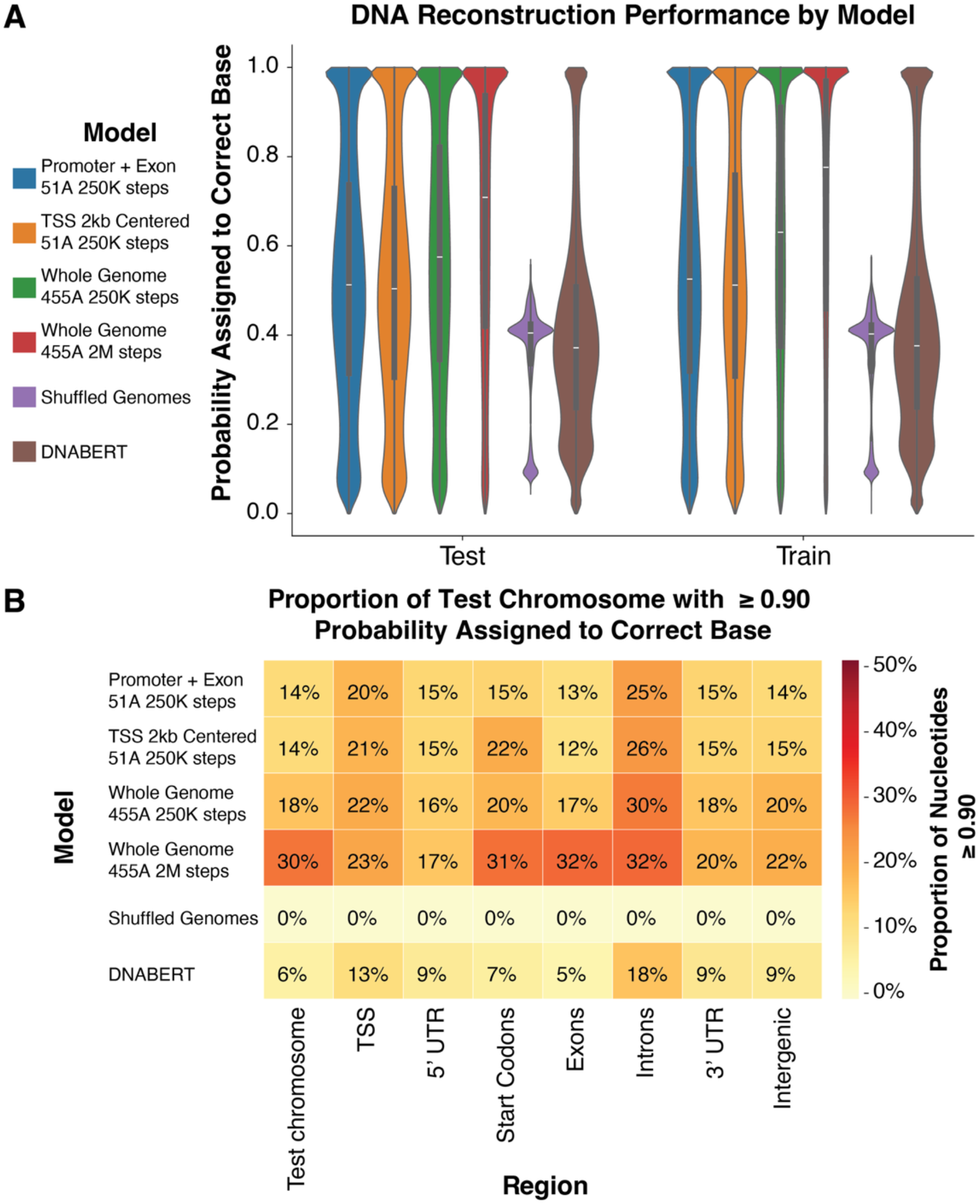
Accurate reconstruction of masked nucleotides in the *P. falciparum* genome. **A)** Violin plots showing the distribution of the probability assigned to the correct base for each DNA reconstruction model. Two annotation-based models are trained on 51 assemblies (“Promoters + Exons 51A 250k steps” and “TSS Centered 51A 250k steps”, respectively). Two models are trained on all 455 assemblies, with one trained for 8 times longer than the other (“Whole Genome 455A 250k steps” and “Whole Genome 455A 2M steps”, respectively). A baseline model is trained on shuffled genomic sequences (“Shuffled Genomes”) and the previously published DNABERT (Ji et al., 2021) model trained on human sequences is shown for comparison. Results for the held-out test *P. falciparum* chromosome 14 are shown on the left, and results for all training chromosomes are shown on the right. **B)** Heatmap showing the percentage of bases in each region where the correct base is assigned a probability of greater than or equal to 0.9 stratified according to genomic annotation. The “Test chromosome” column shows model performance over every base on the test chromosome.

As a negative control, we trained an additional model on the mononucleotide shuffled genomes of all 455 assemblies. This preserved overall genome nucleotide composition but destroyed all small-scale patterns. The “Shuffled Genomes” model produces a distribution of the probability assigned to the correct base centered near 0.4 on both the training set and test set data (**Fig. 2A**). This pattern is consistent with the global A/T content of ∼80% in the *P. falciparum* genome (Gardner et al., 2002). The Shuffled Genomes model does not yield any high-confidence predictions for correct bases, suggesting that the correct high-confidence predictions from trained models reflect learned sequence patterns rather than background nucleotide frequencies.

As an external benchmark, we also evaluated DNABERT, a transformer-based genomic language model pretrained on the human genome (Ji *et al*., 2021). We chose DNABERT as the comparison model because its reconstruction task and prediction outputs are the most comparable to GPN. It was trained on a bidirectional 3-mer token reconstruction task using 512 bp input sequences (see **Methods**). Newer self-supervised models are less directly comparable: DNABERT-2 splits sequence into variable-length tokens, so a masked token does not correspond to a single base (Z. Zhou et al., 2024), and Evo 2 predicts each base only from the sequence upstream of it (Brixi et al., 2026). DNABERT performed worse than all four apicomplexan-specific models on the held-out chromosome. Genome-wide, only 6% of correct test chromosome bases reached a 0.9 high-confidence threshold in the DNABERT model, compared to 14%-30% for the trained GPN models. The median probability assigned by DNABERT to the correct base (∼0.37) was comparable to, and slightly below, the “Shuffled Genomes” control. But unlike the shuffled control, DNABERT produced a tail of high-confidence predictions (**Fig. 2A**). This indicates that DNABERT has learned some transferable sequence features, but that they generalize poorly to the highly A/T biased genome of *P. falciparum*.

Pretraining on phylogenetically relevant sequences is likely driving the improved model performance of the apicomplexan-specific models over DNABERT.

Stratifying correct, high-confidence predictions (defined as a model probability ≥ 0.9 assigned to the correct base) by genomic region reveals several interesting patterns (**Fig. 2B**). We find that introns are the most reliably reconstructed feature across all apicomplexan-specific models, with 25% to 32% of bases in introns correctly predicted with high-confidence. While the “Whole Genome 455A 2M steps” model is the best performing model across all genomic regions (**Fig. 2B**), the benefit of extended training is region-dependent. The “Whole Genome 455A 2M steps” model nearly doubles the fraction of high-confidence correct bases in exons (17% to 32%) and across the test chromosome as a whole (18% to 30%). This model also has a large increase in accuracy near start codons (20% to 31%). However, the longer training time only resulted in marginal accuracy gains in regulatory regions (5’ UTRs: 16% to 17%; 3’ UTRs: 18% to 20%; TSSs: 22% to 23%; and intergenic regions: 20% to 22%). One possible interpretation is that additional training helps the model capture coding-sequence constraints but is not enough to significantly boost performance in less structured and non-repetitive regions like UTRs and intergenic regions. Finally, the DNABERT model falls below all four apicomplexan-specific models in every column, reinforcing the idea that regional gains in the GPN models are specific to apicomplexan pretraining rather than being due to a generic property of genomic language models.

### High reconstruction confidence extends to transcription factor binding sites

Since the apicomplexan-specific GPN model reconstructs a large fraction of the genome with high confidence (**Fig. 2**), we asked whether that reconstruction accuracy highlights regulatory elements, beginning with transcription-factor binding sites. We focused on the *P. falciparum* TF *Pf*AP2-G2, a factor with a well-characterized binding motif and one of the largest sets of ChIP-seq peaks available in *P. falciparum* (6,459 peaks merged from two studies; see **Methods**) (Shang et al., 2022; S. Singh et al., 2021). For each binding site containing the cognate motif (1,841 peaks on training chromosomes, 194 peaks on the test chromosome), we extracted a 200bp window centered on the motif. We then computed the per-base reconstruction probability using the best performing GPN model, “Whole Genome 455A 2M steps” (**Fig. 3**). This model correctly reconstructs most bases around *Pf*AP2-G2 binding motifs with high confidence (**Fig. 3**).

**Figure 3.**
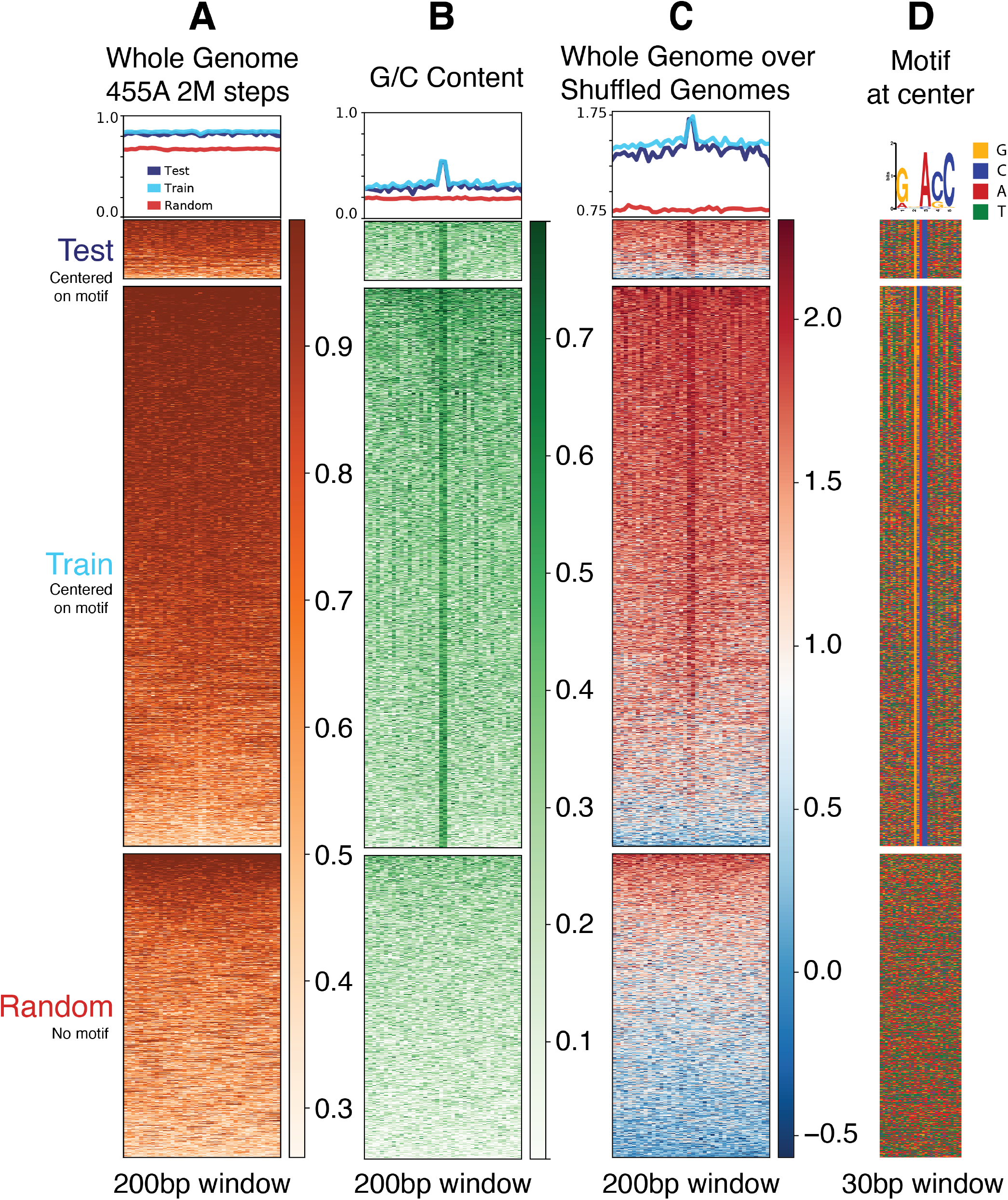
Per-base reconstruction scores and regional context around *Pf*AP2-G2 binding sites. Each row in the heatmaps represents a window centered on a *Pf*AP2-G2 motif occurrence within a ChIP-seq peak. Rows are linked in order across heatmaps and are grouped into three blocks: motif instances in the held-out test chromosome (chr 14; n = 194), instances in the training chromosomes (n = 1,841), and 1,000 random genomic regions sampled from the *P. falciparum* genome. **A)** The per-base probability assigned by the “Whole Genome 455A 2M steps” model to the correct base sorted highest to lowest within each block by the region median score. **B)** Local G/C content across the windows. **C)** The ratio of the scores assigned by the “Whole Genome 455A 2M steps” model to correct bases over the scores from the “Shuffled Genomes” baseline model. **D)** The underlying nucleotide sequence in a narrow 30bp window, also centered on the *Pf*AP2-G2 motif (A is red, T is green, G is orange, and C is blue).

Two patterns emerge. First, reconstruction confidence by the “Whole Genome 455A 2M steps” model at *Pf*AP2-G2 binding sites substantially exceeds the “Shuffled Genomes” baseline. The ratio of scores assigned by the “Whole Genome 455A 2M steps” model to correct bases over the scores from the “Shuffled Genomes” baseline model is consistently elevated across both the test and training blocks and falls towards baseline in random regions. This indicates that the high reconstruction scores reflect learned sequence structure at these sites rather than background A/T base composition. Second, the elevated confidence is a property of the broader peak region rather than the motif itself, indicating that reconstruction probability does not spike at the motif center relative to the surrounding sequence, even when local G/C base content does. The model accurately reconstructs sequences at *Pf*AP2-G2-bound motifs, but it also learns to reconstruct additional sequences in the surrounding 200bp window.

### TF-DNA binding motifs are enriched in highly reconstructible regions

Having shown that reconstruction confidence is elevated across bound *Pf*AP2-G2 motif sites (**Fig. 3**), we asked whether additional regulatory motifs can be identified *de novo* in sequences reconstructed most confidently by the model. We performed motif discovery using XSTREME (Grant & Bailey, 2021) on sets of highly reconstructible sequence windows located within 1 kbp upstream of *P. falciparum* transcription start sites (TSS) (see **Methods**). Highly reconstructible windows covered 9% of the 1 kbp promoter regions. The search yielded nineteen significant DNA sequence motifs enriched in highly reconstructible promoter sequences compared with a control set of random length-matched TSS-proximal sequences (E-value < 0.05).

To determine whether these *de novo* discovered motifs corresponded to known regulatory elements, we compared each against a database of previously characterized *P. falciparum* binding motifs from a *P. falciparum* ApiAP2 protein binding microarray (PBM) study (Campbell et al., 2010) and from ChIP-seq datasets (see **Methods**). We found thirteen of the nineteen motifs have a significant match to an annotated motif (E-value < 1e-3; **Fig. 4**), corresponding to known binding motifs for eight distinct ApiAP2 TFs: *Pf*AP2-I (PF3D7_1007700), *Pf*AP2-L (PF3D7_0730300), *Pf*AP2-G5 (PF3D7_1139300), *Pf*AP2-HC (PF3D7_1456000), PF3D7_1222400, PF3D7_0420300, PF3D7_1107800, and PF3D7_1239200.

**Figure 4.**
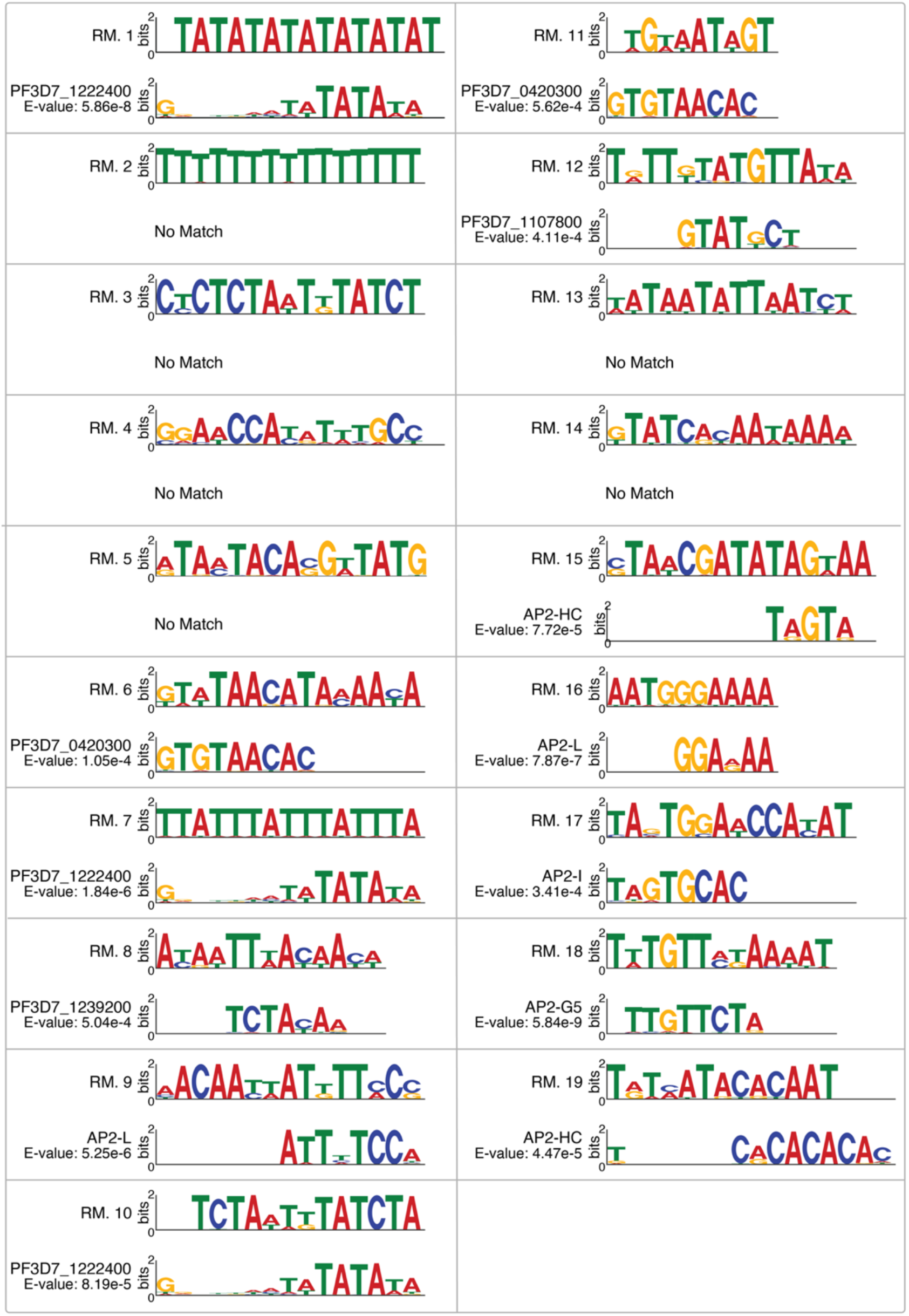
*De novo* motifs discovered in highly reconstructible promoter regions and matches to known *P. falciparum* binding motifs. XSTREME identified nineteen motifs enriched in highly reconstructible windows located within 1 kbp upstream of transcription start sites, relative to length-and number-matched randomly selected non-reconstructible promoter sequences. Each motif is shown as an information-content logo (bits). Motifs are ordered by XSTREME E-value, from most to least significant, proceeding down the left column and then down the right column. In each pair, the top logo is the *de novo* motif recovered from highly reconstructible sequence (RM: “Reconstructed motif”), and the bottom logo is the best-matching previously annotated motif identified by STAMP (max E-value < 6e-4); annotated motifs are drawn from a protein-binding microarray study and from ChIP-seq datasets (see **Methods**). Thirteen of the nineteen motifs had a significant match. The remaining six motifs had no significant database match (“No match”).

Of the six motifs without a significant match to known TF binding motifs, one was a low complexity poly-T motif (RM#2) and another was highly A/T rich (RM#13). The recovery of these motifs is consistent with a fraction of the most reconstructible sequences being repetitive elements rather than cis-regulatory motifs. The four other motifs without known matches may reflect previously unknown regulatory motifs for as-yet unprofiled TFs. Indeed, matching our discovered motifs against sets of motifs previously predicted to be enriched upstream of start codons (Elemento et al., 2007; Young et al., 2008) finds several additional matches (**Supplemental Fig. 3**).

We have shown that reconstruction accuracy is elevated around *Pf*AP2-G2 binding sites (**Fig. 3**), and that highly reconstructible promoter sequences are enriched for known ApiAP2 motifs (**Fig. 4**). Both results indirectly suggest that pretraining allows the model to learn some regulatory motif features from sequence reconstruction alone. To directly test if the model’s internal layers represent known TF binding motifs, we used a neural network interpretability technique called “concept attribution” (Kim et al., 2018) as implemented in the Testing with PCA-projected Concept Activation Vectors (TPCAV) framework (J. Yang & Mahony, 2026).

TPCAV’s concept attribution analysis asks whether a user-defined biological concept is represented within a trained model’s internal activity. Concepts are defined by example; selections of sequences that do or do not represent a given biological concept are presented to the trained model. A linear classifier is then trained to separate concept from non-concept sequences based on the activation pattern of nodes at an intermediate layer of the model. If the classifier can successfully discriminate concept and non-concept sequences, the model can be said to have learned features associated with the concept. TPCAV summarizes concept classification performance with an F-score, the harmonic mean of precision and recall with equal numbers of concept and control examples.

Here, we used TPCAV to evaluate whether the pretrained model encapsulates two categories of genomic concepts. The first category defines DNA motif concepts using 45 known transcription factor binding motifs; concept-representing sequences contain embedded instances of the motif, while non-concept sequences do not (see **Methods**). The second category defines biological annotation concepts using 28 types of genomic regions including coding sequences (CDSs), exons, transcription start sites (TSSs), and 11 chromatin states inferred at 30 hours post-invasion (Brown et al., 2026). In this category, concept-representing sequences are selected based on overlap with the relevant genomic annotation. Based on the average F-scores of the concept classifiers, the GPN model pretrained on randomly shuffled genomic regions displayed the poorest encapsulation of DNA motif concepts (average F-score 0.78 for concept example sequences containing 12 motif insertions, **Supplemental Fig. 4**). The “Whole Genome 455A 2M steps” model performed best in capturing DNA motif concepts (average F-score 0.94 for concept example sequences containing 12 motif insertions, **Supplemental Fig. 4**).

### Fine-tuning strategy affects performance on downstream tasks

Having established that the pretrained model encapsulates regulatory motifs and features, we hypothesized that the pretrained model could be effectively fine-tuned to predict regulatory signals in apicomplexan genomes. To re-use the pretrained model core weights for downstream tasks, we replaced the nucleotide reconstruction output layers with classification layers (blue path in **Fig. 1**). In the following results, each pretrained classifier model is initialized with the model core weights from the “Whole Genome 455A 2M steps” model. Performance will be compared with that of a model with the same architecture but with randomly initialized weights.

As described in the Introduction, the choice of fine-tuning strategy can determine whether the pretraining weights enhance or hinder performance on downstream tasks. In standard fine-tuning, the classification layers are usually initialized at random. Early in training, large gradient updates are required to fit the randomly initialized classification layers, but this will also rapidly alter the model core’s weights if both are optimized simultaneously. This can have the effect of overwriting the features learned during pretraining, and is described by Kumar, *et al*., as the feature distortion effect (Kumar *et al*., 2022). To address this issue, Kumar, *et al*., developed Linear Probe then Fine Tune (LPFT) (Kumar *et al*., 2022) in which the model core weights are frozen at first and only the classifier layers are trained. In later training steps, the model core is unfrozen and the full model is trained end-to-end at a reduced learning rate. This two-phase protocol preserves the pretrained weights during the critical, more volatile early training period and then allows task-specific adaptation once the classifier layers have stabilized.

We tested four pretrained model fine-tuning strategies and compared these against a model with randomly initialized weights (“No Pretraining”). The models were first used to predict binding sites for the *P. falciparum* DNA binding protein *Pf*AP2-I (n = 198 training peaks), and the chromatin-associated heterochromatin protein 1 (*Pf*HP1) (n = 2,931 training peaks) (**Fig. 5**).

**Figure 5.**
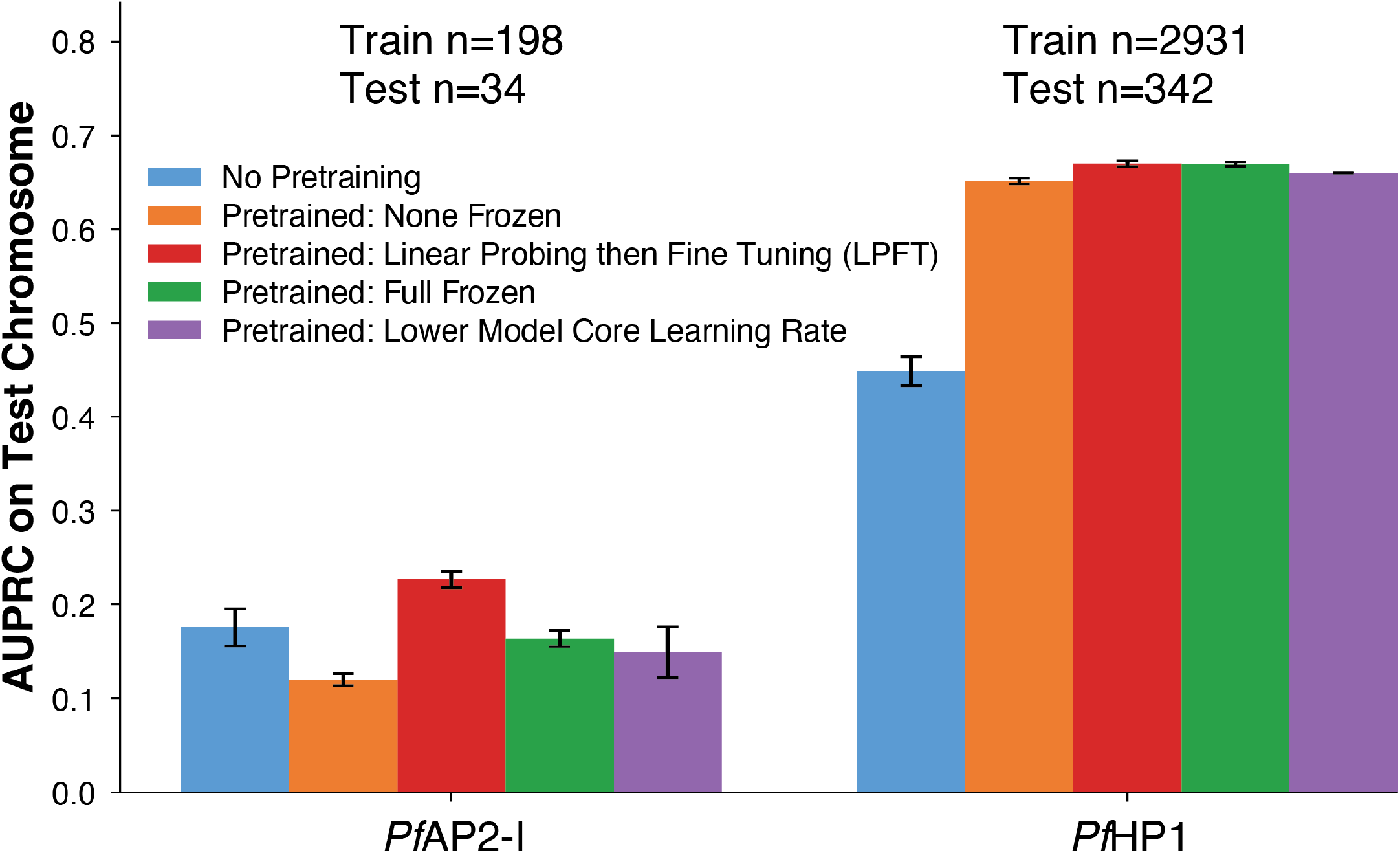
Foundation model fine-tuning strategies for classifiers that predict binding sites of *Pf*AP2-I and *Pf*HP1. Each bar height is the mean of three runs with different seeds, and the bars show one standard deviation above and below the mean. The leftmost bar (blue) is the model without pretraining and each of the other bars represents a different fine-tuning strategy on the pretrained model.

These two datasets were chosen to span a dataset with a limited number of peaks (*Pf*AP2-I) and a dataset with a larger set of available training data (*Pf*HP1). One pretrained model was fine-tuned on the classification task without any weight freezing, allowing all weights to be updated throughout training (“Pretrained: None Frozen”). A second model was trained with the original LPFT strategy by freezing the model core’s weights for half of the training run, then unfreezing and assigning a learning rate 1/10^th^ the rate of the classifier layers (“Pretrained: Linear Probing then Fine Tuning (LPFT)”). Another model was trained keeping the model core fully frozen for the entire training run and only allowing updates to the weights in the classifier layers (“Pretrained: Full Frozen”). Finally, a model was trained with no frozen weights, but with a differential learning rate of 1/10 in the model core compared to the classifier layers (“Pretrained: Lower Model Core Learning Rate”). These were all compared to the randomly initialized model with no pretraining (“No Pretraining”).

With its severely limited training dataset size (198 training peaks), predicting *Pf*AP2-I binding sites demonstrates that the benefit of pretraining critically depends on how the model core weights were updated. The standard end-to-end fine-tuning (“Pretrained: None Frozen”) was counterproductive, as it yielded a mean AUPRC of 0.12 which is 0.06 lower than the model with no pretraining. This is consistent with the feature distortion phenomenon described previously (Kumar *et al*., 2022). In contrast, the LPFT method achieved the highest AUPRC (mean = 0.23), while outperforming the “No Pretraining” model by an AUPRC of 0.05. The “Pretrained: Full Frozen” and “Pretrained: Lower Model Core Learning Rate” strategies showed intermediate performance. Both were better on average than the standard fine-tuning but worse than both the “No Pretraining” model and the “Pretrained: Linear Probing then Fine Tuning LPFT” model.

In the moderate-data target, *Pf*HP1 (2,931 training peaks), all pretrained conditions substantially outperform the model without pretraining, with mean AUPRC scores in the 0.65 to 0.67 range compared to the “No Pretraining” mean AUPRC of 0.45. The gap between pretrained fine-tuning strategies was almost indistinguishable, with the worst strategy (standard fine-tuning) at AUPRC 0.65 and the best (LPFT) at 0.67. This suggests that with sufficient data, the specific training protocol matters less. The absolute AUPRC difference between *Pf*AP2-I and *Pf*HP1 and the order of magnitude difference in data suggests that the benefit of pretraining depends on dataset size while the relative importance of the fine-tuning strategy is greatest when labeled data is scarce. We used LPFT for all subsequent experiments to focus analysis on the relationship between the benefits of pretraining and dataset size.

### Pretraining improves prediction of TF binding sites, chromatin accessibility, and chromatin states

We next sought to determine whether the benefit of pretraining generalizes across a broader range of *P. falciparum* epigenomic signal prediction tasks and whether the benefit depends on the amount of labeled training data. We applied architecture-matched models, with and without pretraining, to predict *P. falciparum* binding sites of fourteen ApiAP2 transcription factors (*Pf*AP2-EXP, *Pf*AP2-G2, *Pf*AP2-G3, *Pf*AP2-G5, *Pf*AP2-G, *Pf*AP2-HC, *Pf*AP2-I, *Pf*AP2-LT, *Pf*AP2-L, *Pf*AP2-O2, *Pf*AP2-O5, PF3D7_1239200, PF3D7_0613800 and *Pf*AP2-P) and two chromatin-associated factors (MORC and *Pf*HP1) (see **Methods**). These selected factors span roughly two orders of magnitude in available ChIP-seq peaks in the training chromosomes (n = 198 to n = 10,310). We evaluated model performance by area under the precision-recall curve (AUPRC) on sequences drawn from the entire test chromosome (chr 14) (**Fig. 6A**).

**Figure 6.**
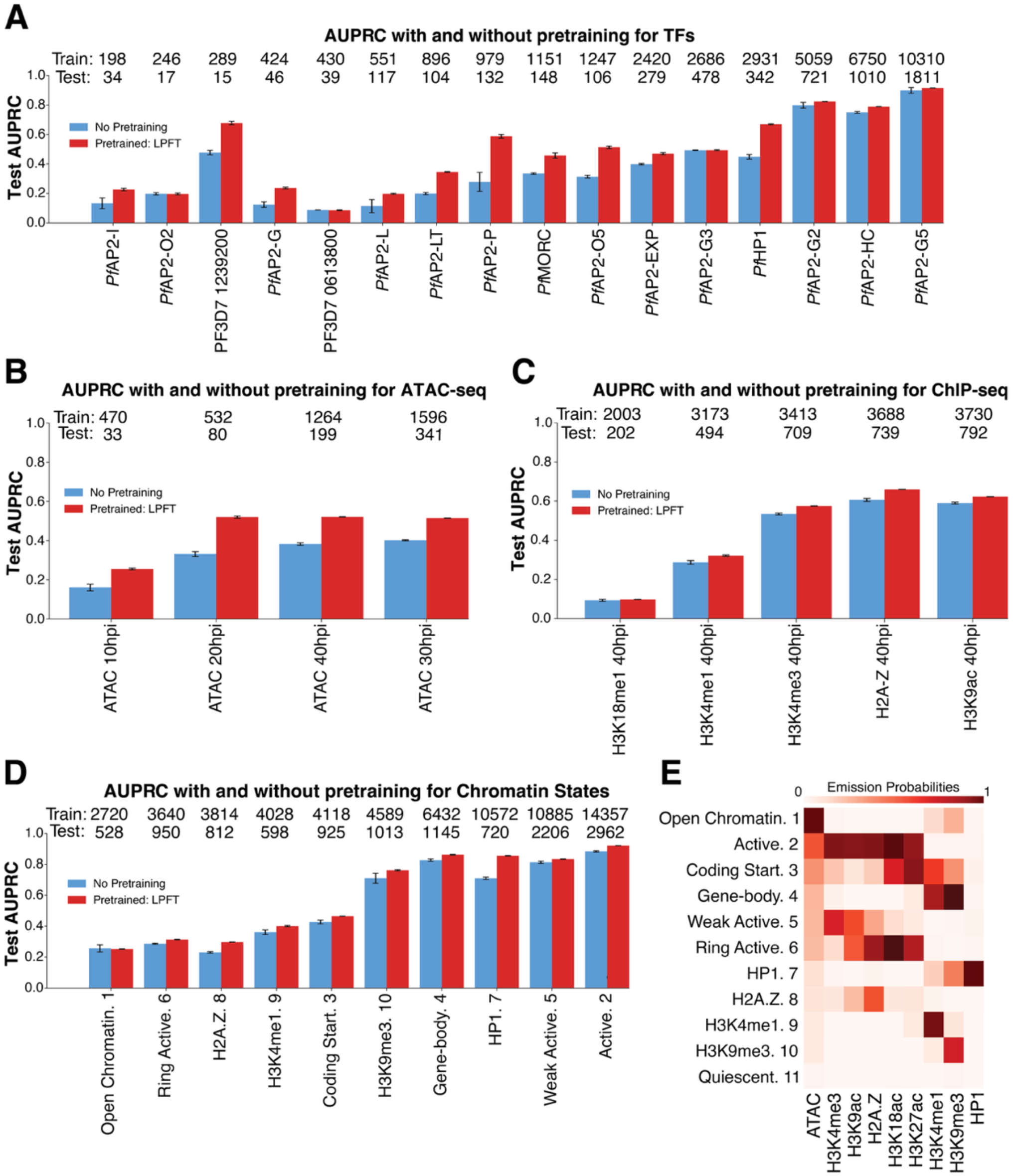
AUPRC on held-out test chromosome 14 for various epigenomic classification models, without pretraining (blue) and with foundation model-based LPFT fine-tuning (red). **A)** AUPRC for predicting *P. falciparum* binding sites of fourteen ApiAP2 TFs and two chromatin-associated factors (MORC and *Pf*HP1). Factors are ordered left to right by increasing number of ChIP-seq peaks in the training datasets (the number of training and test peaks are above each pair of bars, training on top, then test on bottom). Error bars represent ± 1 standard deviation across three random seeds. **B)** As in (A), for classifiers predicting *P. falciparum* peaks of chromatin accessibility measured by ATAC-seq at four time points spanning the asexual developmental cycle (10, 20, 30, and 40 hours post-invasion; hpi). **C)** As in (A), for classifiers predicting *P. falciparum* peaks of four histone modifications and one histone variant measured by ChIP-seq at 40 hpi. **D)** As in (A), for classifiers predicting regions displaying ten of the eleven *P. falciparum* ChromHMM chromatin states defined in Brown, et al. (2026). The quiescent state (state 11) was excluded from direct evaluation because it represents a lack of enriched epigenomic signals and is highly abundant across the genome. States are ordered by increasing number of 200 bp genomic bins assigned to each state in the training datasets. State names and numerical identifiers follow Brown, *et al*. (2026). **E)** ChromHMM hidden Markov model emission probabilities for all eleven chromatin states across nine chromatin features (ATAC-seq, H3K4me3, H3K9ac, H2A.Z, H3K18ac, H3K27ac, H3K4me1, H3K9me3, and *Pf*HP1 ChIP-seq), reproduced from Figure 1A of Brown, *et al*. (2026). Color intensity indicates the probability of observing each chromatin signal within a genomic bin assigned to each state.

Pretraining improved TF binding predictions for most factors, but the magnitude of the benefit was strongly influenced by the size of the training dataset. The largest and most consistent gains were for datasets with fewer than 3,000 training peaks. The median increase in AUPRC was 0.09 across the full panel, but the median AUPRC improvement increases to 0.11 among factors with fewer than 3,000 training peaks. The largest gains appeared in the ApiAP2 TF *Pf*AP2-P (delta AUPRC = 0.31, n = 979 training peaks) and the heterochromatin-associated factor *Pf*HP1 (delta AUPRC = 0.22, n = 2,931 training peaks). Both factors bind in sub-telomeric heterochromatic regions which often contain repetitive sequences that are likely to be well represented in the reconstruction model weights. In contrast, the three factors with more than 3,000 training peaks (*Pf*AP2-G2, *Pf*AP2-HC, and *Pf*AP2-G5) and three of the factors with fewer than 3000 training peaks (*Pf*AP2-O2, PF3D7_0613800, and *Pf*AP2-G3) showed marginal or no benefit from pretraining. This indicates that the marginal value of pretrained representations diminishes once enough labeled data are available for the downstream model to learn comparable features. The very-low-data examples (fewer than 500 training peaks) tended to have low overall AUPRC scores and less benefit from pretraining. The striking exception is PF3D7_1239200 with only 289 training peaks but with an AUPRC increase of 0.2, which shows pretraining can be exceptionally beneficial in some small datasets. PF3D7_1239200 binding sites are mostly found in highly repetitive sub-telomeric regions (Shang et al., 2022), which, due to their repetitive nature, are well represented in the pretrained model’s weights. Our findings mirror previous reports that pretraining gains are largest in low-data settings (Novakovsky et al., 2021; Z. Tang et al., 2025).

To assess whether our GPN model and pretraining strategy offer beneficial performance compared with a previous application of deep learning to *P. falciparum*, we reimplemented the convolutional–recurrent architecture of MalariaSED (Wang et al., 2023). MalariaSED is a previous application of deep learning models to predict epigenomic features in apicomplexan genomes, which demonstrated that neural networks can predict epigenomic signature changes resulting from strain-specific mutations in *Plasmodium falciparum*. We compared our MalariaSED-like model’s performance with that of our pretrained GPN classifiers across sixteen DNA-binding factors (**Supplemental Fig. 5**). Pretrained GPN classifiers significantly outperformed the MalariaSED-like models on ten of sixteen factors and performed comparably on four, confirming the benefits of pretraining on DNA-binding factor prediction tasks.

We next determined whether the benefits of pretraining extend to predicting chromatin accessibility (measured by ATAC-seq) and histone modifications (measured by ChIP-seq) in *P. falciparum* (**Fig. 6B-C**). Across four ATAC-seq timepoints (n = 470 to 1,596 training peaks) spanning the asexual blood stages of parasite development, pretraining consistently improved prediction of chromatin accessibility (median AUPRC increase = 0.13, range: 0.09 to 0.19; **Fig. 6B**). This is comparable to the stronger TF gains with fewer than 3,000 peaks. The histone modification ChIP-seq datasets had higher numbers of peaks (n = 2,003 to 3,730 training peaks) and generally smaller performance increases than the ATAC-seq data (median AUPRC increase = 0.03, range: 0.004 to 0.053; **Fig. 6C**), consistent with our observation that pretraining provides diminishing returns with increasing training data. One target, H3K18me1 (n = 2,003 training peaks), was poorly predicted regardless of pretraining (AUPRC ∼ 0.1 for both models), suggesting that this dataset is an outlier and is largely unpredictable from local sequence.

H3K18me1 is anti-correlated with active marks, is not strongly associated with gene expression, and is found primarily near centromeres (Reers et al., 2023). Both centromeres and sub-telomeric regions are characterized by simple repeats, but the distribution of H3K18me1 favors centromeres. The model’s poor predictive performance on H3K18me1 suggests that sequence context alone cannot differentiate these types of repetitive regions, resulting in limited predictive performance.

Finally, we evaluated predictions on composite chromatin states, which integrate accessibility, histone modifications, and *Pf*HP1 occupancy into discrete regulatory categories. We used our previously published eleven-state ChromHMM annotation defined for *P. falciparum* and predicted ten of the eleven states (only excluding the “quiescent” state which is the most common state and covers ∼30% of the genome) (**Fig. 6D**) (Brown et al., 2026). The emission probabilities defining each state are reproduced in **Fig. 6E** based on Figure 1 in Brown, *et al*. (2026). There are comparatively high numbers of regions displaying each state on the genome (n = 2,720 to 14,357 training regions) because the genome was segmented into 200bp bins with one chromatin state per bin. Pretraining produced small gains overall (median AUPRC increase = 0.04) with the exception of state 7 (“HP1. 7”) which has an AUPRC increase of 0.15 (n = 10,572 training regions). The large benefit of pretraining for state 7, despite its relatively high number of observations on the test chromosome, mirrors the large AUPRC gain for the transcription factor ChIP-seq dataset *Pf*HP1 (AUPRC increase for *Pf*HP1 = 0.22) (**Fig. 6A**). This suggests that clade-specific pretraining can capture repetitive features and other distinctive sequence composition in *P. falciparum* heterochromatic regions (Flueck et al., 2009; Gardner et al., 2002; Hernandez-Rivas et al., 2010).

### Truncated foundation models retain pretraining advantages on downstream classification tasks

While we have demonstrated that pretrained GPN models offer advantageous classification performance on downstream tasks with limited training data, networks that use the entire 25 convolutional layer GPN model core likely have excessive numbers of parameters for most downstream tasks and are costly to fine-tune. It is possible that the most informative features for downstream epigenomic tasks are encoded in the initial convolutional layers of the GPN, and therefore only a subset of layers may be necessary to gain the advantages of pretraining. To test this hypothesis, we truncated the 25-layer GPN model core (“Whole Genome 455A 2M steps” model) to its first *N* convolutional layers (*N* = 2, 4, 8, 16). We compared performance on downstream TF binding site prediction tasks after LPFT fine-tuning. We evaluated predictions in six TF binding datasets, spanning the available range of training set peak counts (*Pf*AP2-I, n = 198; *Pf*AP2-G, n = 424; *Pf*AP2-P, n = 979; *Pf*HP1, n = 2,931; *Pf*AP2-G2, n = 5,059; *Pf*AP2-G5, n = 10,310), and report mean test-chromosome AUPRC across three random seeds (**Fig. 7**).

**Figure 7.**
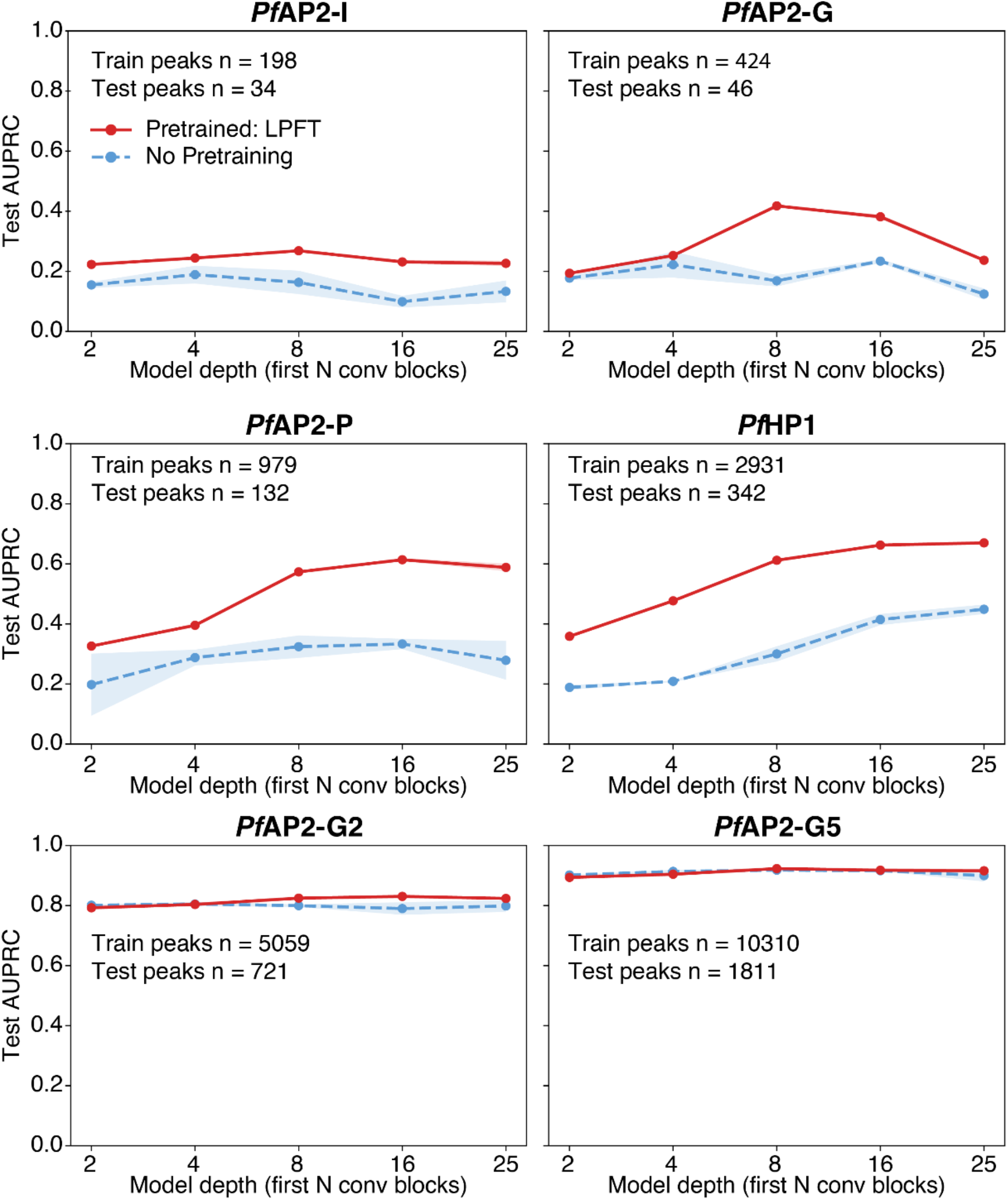
Classification performance of GPN networks with and without pretraining as a function of model core depth for six *P. falciparum* DNA-binding factors. For each factor, the 25-layer GPN model core was truncated to its first N convolutional layers (N = 2, 4, 8, 16) and compared to the full 25-layer model. For each factor we compared a model with no pretraining (dashed blue lines) to a pretrained LPFT fine-tuned model (solid red lines). Each point is the mean AUPRC on the held-out chromosome (chr 14) across three random seeds; shaded bands show one standard deviation around the mean. Panels are arranged from left to right, top to bottom in order of increasing test-chromosome peak count.

For the most data-limited factors, the pretraining advantage was concentrated in the earliest layers. With only 2 convolutional layers, the models without pretraining had very poor performance (AUPRC = 0.15 for *Pf*AP2-I and 0.18 for *Pf*AP2-G), while the pretrained models using the first two layers of the foundation model already reached AUPRC = 0.22 and 0.19, respectively. For both factors, the pretraining performance peaked at 8 layers (AUPRC = 0.27 for *Pf*AP2-I and 0.42 for *Pf*AP2-G). This indicates that at least some relevant sequence features are already encoded in the first few layers of the pretrained representation. As model depth increases, the non-pretrained models improve and the gap narrows, but with different patterns for the two cases. For *Pf*AP2-I, both the pretrained model performance and the model without pretraining performance were essentially flat across depth (pretrained AUPRC = 0.22 at two layers, a shallow peak of 0.27 at eight layers, and 0.23 at full depth). The model without pretraining peaked at 4 layers (AUPRC = 0.19) where it had the most similar performance to the pretrained model (AUPRC difference = 0.06). The benefit of pretraining increased slightly from an AUPRC improvement of 0.07 at two layers to 0.09 at the full depth model. For *Pf*AP2-G, the pretrained model peaked at eight layers (AUPRC = 0.42, an increase of 0.25 compared to the model without pretraining) and then declined, converging with the model without pretraining by the full 25-layer model (with an AUPRC difference of 0.11); the pretraining benefit for *Pf*AP2-G was almost entirely lost at full depth.

For the factors with fewer than 3,000 peaks, where pretraining provides its largest and most consistent benefit (**Fig. 6A**), the advantage of pretraining is also realized early but it persists across depths. The pretrained models reached roughly 90% of their eventual performance by eight layers. For example, predicting *Pf*AP2-P peaks has an AUPRC = 0.57 at eight layers versus AUPRC = 0.61 at sixteen layers. Predicting *Pf*HP1 peaks has an AUPRC = 0.61 at eight layers versus AUPRC = 0.67 at the full depth. The models without pretraining showed a more consistent increase in performance as the model deepened but they did not close the gap with the pretrained model. Here, additional depth modestly helps performance of the pretrained model up to roughly eight to sixteen layers, but the bulk of the benefit was again captured by the first third of the network.

For the two data-rich classification tasks, depth and pretraining were both nearly irrelevant. Performance when classifying *Pf*AP2-G2 and *Pf*AP2-G5 peaks is already high using two GPN layers (AUPRCs: *Pf*AP2-G2 = 0.79, *Pf*AP2-G5 = 0.89) and was practically flat across different depths. The model without pretraining performed nearly as well as the pretrained model, within 0.04 AUPRC for *Pf*AP2-G2 and was essentially indistinguishable for *Pf*AP2-G5 (gap ≈ 0.00– 0.02). With abundant training data labels, the naïve model learns comparable features at shallow depth. Therefore, neither extra depth nor pretrained initialization adds to the performance meaningfully. This is consistent with the diminishing returns we observe across the full panel (**Fig. 6A**).

## Discussion

Predictive deep learning models promise several compelling applications in regulatory genomics, including for the genomes of eukaryotic parasites such as *Plasmodium spp*. However, training neural networks for application to small genomes is inherently constrained by the limited sizes of training datasets compared with larger eukaryotic genomes. Attention has recently focused on using self-supervised foundation models to initialize regulatory genomic classifiers, including for small eukaryotic genomes (Karollus et al., 2024). Our findings support clade-specific foundation model pretraining as a practical route to data-efficient regulatory modeling in small eukaryotic genomes. Across varied predictive tasks, including classifying TF DNA-binding sites, histone marks, accessible regions, and chromatin states, pretraining improved binding prediction for most targets. However, the magnitude of improvement provided by pretraining was strongly influenced by training label abundance (**Fig. 6A**). Performance gains were largest and most consistent for transcription factors with fewer than 3,000 ChIP-seq peaks in the training set, while the improvement shrank to near zero once a factor had more than 3,000 training peaks available. This pattern is important in the face of the main critique regarding genomic foundation models; i.e., that carefully trained task-specific models can match or exceed pretrained models on data-rich benchmarks (Feng et al., 2025; Z. Tang et al., 2025; You et al., 2025). Our results suggest that this assessment and the case for pretraining are not in conflict. Rather, the value of pretraining is conditional on data scarcity.

Our interpretation of this result is that self-supervision on hundreds of related genomes front-loads regulatory sequence structure into the model core’s weights. Models with randomly initialized weights must learn relevant features from a handful of TF-DNA binding sites without prior knowledge of common patterns. Once sufficient labeled data are available, task-specific models can recover comparable features on their own and the advantage of pretraining disappears. This diminishing returns result is consistent with prior reports that pretraining gains are largest in low-data settings (Novakovsky et al., 2021; Z. Tang et al., 2025).

We also note that the benefit we observe is not a generic property of large pretrained models but depends on the phylogenetic proximity between the pretraining and target sequences.

DNABERT, a transformer language model pretrained on the human genome (Ji et al., 2021), transferred poorly to *P. falciparum* (**Fig. 2A**). This is likely due to the additional problems posed by sequence composition and distinct regulatory architecture. The *P. falciparum* genome is highly A/T rich (80% A/T on average) and has many small tandem repeats (Gardner et al., 2002). Sequences with regulatory function must therefore be distinguishable from a genome with very different properties than most eukaryotic genomes. Pretraining within the clade addresses this obstacle by training on genome assemblies that are more similar to *P. falciparum*. Similar results have been reported in plants: family-level models of histone modifications transfer substantially better within *Poaceae* or *Brassicaceae* than across monocot–dicot boundaries (Lv et al., 2026), and a foundation model pretrained specifically on angiosperm genomes (PlantCAD2) outperforms much larger generalist models on plant regulatory tasks (Zhai et al., 2026).

The strategy used to adapt the pretrained model core was most consequential in the cases where pretraining mattered most. For severely data-limited tasks, standard end-to-end fine-tuning was counterproductive, underperforming the model without pretraining. This is consistent with the feature-distortion phenomenon in which large early gradient updates overwrite useful pretrained representations (Kumar et al., 2022). Linear-probe-then-fine-tune (LPFT), which freezes the model core until the classification layers stabilize before adapting at a reduced learning rate, instead preserved and exploited the pretrained features and achieved the best performance (**Fig. 5**). The practical implication is that in the low-data regime where pretraining delivers the largest benefit, the fine-tuning protocol is also most likely to determine whether those benefits are realized, and a conservative freeze-then-adapt schedule is the safer default.

Our experiments also demonstrated that pretraining smaller network architectures with truncated foundation models can still be effective in low-data settings, offering a path to gaining the benefits of pretraining in models that can be trained more quickly and that are less likely to be subject to over-parameterization. We observed that the benefit of pretraining was substantially realized within roughly the first eight of twenty-five convolutional layers (**Fig. 7**).

This is consistent with TF binding site prediction being a local, motif-scale prediction problem; early convolutional layers may act as motif scanners that capture the relevant short-range structure, while deeper layers encode longer-range, reconstruction-specific features that do not transfer well to the binding task and can even interfere with it. A shallow slice of the pretrained model core is therefore sufficient for downstream binding prediction and, for the most data-scarce factors, could be preferable to the full-depth model.

Beyond the value of our apicomplexan-specific foundation model for pretraining classification models, the variable ability of our model to accurately reconstruct nucleotides along apicomplexan genomes may serve as a useful indicator of functional elements. Motif discovery in the most reconstructible promoter windows recovered a mixture of motifs, several of which matched known TF binding motifs in *P. falciparum* (**Fig. 4**). We acknowledge that some of the discovered motifs could be false positives from XSTREME. However, given the high percentage of matches to known motifs, we believe reconstruction confidence could serve as an annotation-free prior for locating candidate regulatory elements.

Our investigation of clade-specific foundation models is subject to several limitations. Our evaluation rests on a single held-out species — *P. falciparum* — due to limited regulatory genomics data existing in other apicomplexan species. Therefore, the extent to which the pretrained representation transfers to other apicomplexan genera such as *Toxoplasma*, *Cryptosporidium*, or *Theileria* remains to be established. Training dataset size is an imperfect proxy for the ease of a prediction task. Some targets, such as H3K18me1, were poorly predicted regardless of pretraining (AUPRC ≈ 0.1 for both models), indicating a ceiling on what local sequence can explain for certain epigenomic signals. Finally, the GPN architecture has limited capacity for long-range modeling relative to more recent designs; a more expressive model core or a wider receptive field might learn comparable features from less data and thereby narrow the pretraining gap, as has been observed in data-rich genomic settings (Avsec et al., 2021, 2026).

Parallel efforts in plants have likewise found that phylogenetically focused pretraining or family-level training improves regulatory prediction relative to broader models (Lv et al., 2026; Zhai et al., 2026), suggesting the strategy may generalize to other data-sparse clades. Our work shows clade-specific foundation models are a useful strategy for pretraining classification models in Apicomplexa, a clade of eukaryotic organisms that possess relatively small genomes and several unusual regulatory features. Because the gains we observe may depend on phylogenetic proximity between sequences in the pretraining and target sets, extending this approach will likely require making separate models for each target clade. Future work should investigate the benefits of self-supervised foundation models in additional clades where small genome sizes and a scarcity of experimental annotation currently constrains the application of deep learning models.

## Methods

### Model training

The GPN-based reconstruction models were trained on DNA sequence from 455 apicomplexan assemblies downloaded from the National Center for Biotechnology Information (NCBI) (Sayers et al., 2025). Two chromosomes from *P. falciparum* 3D7 were withheld from training: chromosome 13 (Pf3D7_13_v3) was used for the validation set because it is the second largest chromosome, and chromosome 14 (Pf3D7_14_v3) was used for the test set because it is the largest chromosome. To prevent leakage of sequences homologous to the held-out chromosomes, we created a per-assembly blacklist based on the held-out chromosomes. To create the blacklists, each training assembly was aligned to the two held-out chromosomes with Minimap2 (v2.28) (H. Li, 2018) using the full-genome alignment option “asm5.” The resulting alignments were sorted with SAMtools (v1.21) (H. Li et al., 2009), converted to BED intervals and merged with bedtools and bamtobed (v2.31.1) (Quinlan & Hall, 2010).

Each of the four reconstruction models used different subsets of regions and assemblies during dataset construction. Only 51 of the assemblies had associated annotation files and these annotated assemblies were used for the “Promoters+Exons 51A” and the “TSS Centered 51A” models. For the annotation-dependent models, we defined a whitelist of regions based on a 1 kb window upstream of the transcription start site (TSS) plus all exons for one model (“Promoters+Exons 51A”) and a whitelist of 2 kb windows centered on the TSS for the other (“TSS Centered 51A”). TSSs were defined based on the 5’ untranslated regions (5’ UTR) annotated for each transcript in the accompanying gene transfer format (GTF) file. After excluding the blacklist, the whitelist regions for each model were binned into 512 bp bins in a sliding window of 256 bp (50% overlap). Both the forward and reverse strand sequences for each bin were included in the dataset for training. Any window with an ambiguous “N” nucleotide was removed. For the two annotation-independent models (“Whole Genome 455A” and “Whole Genome 455A 2M steps”), all 455 assemblies were used, and each assembly was binned into 512 bp windows with a 256 bp offset after excluding the blacklist (as in the annotation-dependent models). The “Whole Genome 455A 2M steps” model trained for the first 200,000 steps on the same windows as the “Whole Genome 455A” model. For each subsequent block of 200,000 steps, the “Whole Genome 455A 2M steps” model’s training data received additional shuffling. Every 200,000 steps after the first 200,000, a random seed was used to randomly shift all of the training windows by some integer in the range [-128, 128]. During training, for all four models, 15% of each 512 bp sequence (∼ 77 bp) was masked at random for the model to predict.

All four models were implemented in PyTorch (version 2.5.1) and wrapped with HuggingFace libraries. The models are based on the Genomic Pretrained Network (GPN) architecture (Benegas et al., 2023). The GPN consists of a dilated CNN model core with 25 convolutional layers with a dilation schedule that doubles between each layer for 6 layers, then is set back to 1 (no dilation), this occurs three times with a final layer with no dilation at the end. The kernel size is 9 for all layers and the hidden size (the number of output channels) is 512 for all layers. Each layer consists of two residual sub-blocks applied sequentially. The first is the 1D convolution with padding set to “same” to preserve sequence length, followed by a GELU and layer normalization. The output of this block is added to the block’s input via a residual connection. The second block is a feedforward block: linear projection of 512 to 512, followed by GELU and layer normalization, with the output again added back via a residual connection. The output of the final convolutional layer is passed to the reconstruction layers. The reconstruction layers are a linear projection of 512 to 512, followed by a GELU, layer normalization, and a final linear projection to the vocabulary size producing per-position logits over the vocabulary. These logits are compared against the true nucleotide identities at masked positions using cross-entropy loss.

The four reconstruction models used AdamW optimizer with a learning rate of 1e-4, weight decay of 0.01, and a cosine learning rate schedule with 1,000 warmup steps. They were trained using mixed-precision (fp16) and a batch size of 512. Models were trained for 250,000 steps with the exception of “Whole Genome 455A 2M steps” which trained for two million steps.

Checkpoints were evaluated every 1,000 steps and the best-performing checkpoint by evaluation loss was retained at the end of training.

As baselines for reconstruction, we compared the GPN models to a pretrained DNABERT (Ji et al., 2021) model and to a GPN model trained on shuffled genomes. For DNABERT, we used the publicly available model on HuggingFace (ID: “zhihan1996/DNA_bert_3”) without fine-tuning it on apicomplexan sequence. Per-base reconstruction probabilities were computed on the *P. falciparum* 3D7 reference genome in sliding 512 bp windows with 250 bp overlap between consecutive windows. Because DNABERT represents sequences as overlapping 3-mers, masking only the central 3-mer on a target base would leak that base through neighboring tokens because the 3-mers have a step size of one. Therefore, for each target position p, we simultaneously masked the three contiguous 3-mer tokens that contain p (p -1, p, and p + 1).

Within each window, multiple target positions were scored in the same forward pass by masking disjoint triples separated by a stride of 50 bp. Softmax probabilities over the 3-mer vocabulary were marginalized over the 16 possible tokens with the desired center base to obtain a renormalized distribution over A, C, G, and T. Retaining the probability of the reference base gave us the reconstruction score.

The “Shuffled Genomes” model used the same GPN architecture but was trained on sequence in which the nucleotide order was destroyed while composition was preserved. Each of the 455 assemblies was cleaned to only contain A, C, G, or T nucleotides (removing any ambiguous “N” bases). Then each assembly was randomly shuffled so that its overall composition was retained but any information content was lost. The shuffled assembly sequences were divided into training, validation, and test with the ratio of 7:1.5:1.5. This control model was trained for 20,000 steps with the same parameters as the other GPN models.

### Reconstruction scores at *Pf*AP2-G2 motif-containing peaks

AP2-G2 ChIP-seq raw fastq files were sourced from two studies (Shang et al., 2022; S. Singh et al., 2021). Reads were aligned with BWA MEM (v0.7.17) with the -M option (H. Li, 2013). Mapped reads were filtered to include only reads with a phred score of at least 30 with SAMtools (v1.16.1) (H. Li et al., 2009). Duplicate reads were removed with Picard Tools’ MarkDuplicates using the parameters REMOVE_DUPLICATES=true and VALIDATION_STRINGENCY=STRICT (The Broad Institute, 2021). We merged replicates after removing duplicates and called peaks using MACS2 (v2.2.2.1) (Zhang et al., 2008) with the options: -g 2e+7 -q 0.001 --nomodel --shift 0 --extsize 200. Summits from the four samples (10, 25, and 40 hours post invasion from Shang et al., (2022) and 30 hours post invasion from Singh et al., (2021)) were pooled and merged to make a single set of non-overlapping peaks with BEDTools. *De novo* motif discovery was performed on the merged regions with MEME-ChIP from the MEME Suite (v5.5.1) (Bailey et al., 2015; Machanick & Bailey, 2011). Peaks containing the AP2-G2 cognate motif were identified via FIMO: 1,841 on the training chromosomes and 194 on the held-out test chromosome.

Region centering, matrix computation, and plotting were carried out using deepTools (v3.5.1) (Ramírez et al., 2016), BEDtools (v2.30.0) and bedGraphToBigWig (v357) from UCSC (Kent et al., 2010). The motif .bed file was split into two sets of regions: all regions on the training chromosomes, and all regions on the test chromosome. We centered each motif and expanded a window around the motif midpoint to 200 bp (100 bp on each side) with computeMatrix reference-point center at 5 bp resolution with deepTools. As a negative control, the genome was tiled into overlapping 200 bp windows (100 bp step size) with bedtools makewindows. We removed any control windows overlapping the 200 bp motif windows and then sampled 1,000 bins from the remaining set. The three region groups: test chromosome motifs, training chromosome motifs, and random windows, are shown in separate panels in (**Fig. 3**). We ordered rows based on the median signal reconstruction score from the “Whole Genome 455A 2M steps” model. This ordering was maintained for all other tracks so that a given row corresponds to the same motif window across panels. Heatmaps and profile plots were drawn with plotHeatmap and plotProfile respectively from deepTools. Local G/C content was computed across the genome with bedtools nuc and used to plot the “G/C Content” heatmap and profile plot in the same motif windows. The ratio of the reconstruction scores from the “Whole Genome 455A 2M steps” model to those from the “Shuffled Genomes” model were used to plot the “Whole Genome over Shuffled Genomes” column (**Fig. 3C**). The rightmost column in the figure (**Fig. 3D**) shows the underlying nucleotide sequence in a narrow 30 bp window centered on the motif.

### Motif discovery in highly reconstructible regions

Per-base reconstruction model scores (i.e., probability assigned to correct base) were smoothened across 5 bp windows with a 1 bp step size. Sequence windows in the top 5% (95^th^ percentile) of scores are defined as highly reconstructible. Overlapping highly reconstructible windows were merged and then expanded to a minimum window size of 30 bp by symmetrically extending both sides of the window. Windows whose coordinates would have extended past either end of a chromosome boundary were truncated. To bound the maximum length of any single region for downstream motif analysis, every merged interval longer than 100 bp was recursively bisected at its midpoint until no piece exceeded 100 bp. No bases were dropped during the bisection and odd numbers of bases were split with the left half receiving 1 more base than the right half.

To focus the analysis on putative regulatory sequences, we restricted the highly reconstructible windows to those where at least half the length of the window overlapped 1 kbp regions immediately upstream of annotated transcription start sites (TSSs) in the *P. falciparum* 3D7 genome. We created a length and number matched set of random intervals from within the 1kb promoter regions to act as a control for the motif search step. We used XSTREME from the MEME suite (v5.5.1) (Bailey et al., 2015; Grant & Bailey, 2021) to search for motifs in the highly reconstructible regions within promoters.

To determine whether these *de novo* motifs corresponded to known regulatory elements, each XSTREME-derived motif was compared against a reference database of previously characterized *P. falciparum* motifs using STAMP (v2.0) (Mahony & Benos, 2007). STAMP aligns each query motif against every motif in the supplied database, scores the alignments with a column-comparison metric, and ranks the database motifs by the E-value of the alignment. We took the top-ranked motif as the match for each query if the STAMP E-value was less than 1e-3. The reference database contained known ApiAP2 motifs from a protein-binding microarray study (Campbell et al., 2010) and motifs obtained by running MEME on several ChIP-seq datasets (see **Datasets** below). We used the default STAMP options of Pearson Correlation Coefficient for the column comparison metric and Ungapped Smith-Waterman for the alignment.

### Downstream classification tasks

We trained binary sequence classifiers for fourteen ApiAP2 TFs (*Pf*AP2-I, *Pf*AP2-O2, PF3D7_1239200, *Pf*AP2-G, PF3D7_0613800, *Pf*AP2-L, *Pf*AP2-LT, *Pf*AP2-P, *Pf*AP2-O5, *Pf*AP2-EXP, *Pf*AP2-G3, *Pf*AP2-G2, *Pf*AP2-HC, and *Pf*AP2-G5), two chromatin-associated factors (MORC and *Pf*HP1), four ATAC-seq timepoints (10, 20, 30, and 40 hours post invasion), four chromatin marks (H3K18me1, H3K4me1, H3K4me3, and H3K9ac), one histone variant (H2A.Z), and ten chromatin states (states 1 through 10 from Brown et al., (2026)) (see **Datasets**).

For each target, we built a training dataset with 512 bp windows. Positive examples were generated by sampling summit-centered windows with a random shift that kept the summit at least 40 bp from either edge of the window. We augmented each target to have 200,000 positive windows. Where it was impossible to create 200,000 unique windows we made all possible unique windows given the constraints above. Negative examples were sampled as a one-to-one mixture of genome-wide random samples and G/C matched samples (±0.05 G/C content). We excluded any window from the negative set if it overlapped a 756 bp buffer window upstream and downstream of a peak summit. The validation and test chromosomes were the same held-out chromosomes used in the reconstruction model (chromosome 13 and chromosome 14, respectively). Validation and test sets used exhaustive chromosome-wide construction; negatives were selected from all genomic windows not in a peak or flanking a peak (within 756 bp of a summit) and positives came from every peak-centered 512 bp window. Both negative and positive windows were sampled so that on average there were five windows selected for every peak summit and five windows overlapping every possible negative base.

Each classifier reused the convolutional GPN model core, replacing only the reconstruction layers with classification layers. The pretrained and “No Pretraining” classifiers shared this architecture and differed only in how the model core was initialized. For the pretrained classifiers, model core weights from the reconstruction task were used as the initial weights in the model core for the classifier task. Because the reconstruction layers and the classification layers are architecturally different (see **Model training**) the weights in the classification layers had to be randomly initialized. For the classifier models without pretraining, the same model core architecture was instead randomly initialized, so no information from the reconstruction task was carried over. The classification layers average the final convolutional hidden states across the 512 bp input sequence, collapsing the per-position 512-dimensional features into a single 512-dimensional vector. That pooled representation is passed through dropout (probability 0.1), a linear projection of 512 to 512, a GELU nonlinearity, a second dropout (probability 0.1), and a final linear projection to two class logits (bound vs unbound). The resulting sequence-level logits were trained with cross-entropy against the binary label of the sequence. To assess how much of the pretrained representation the task requires, the model core was truncated to its first N convolutional layers (N ∈ {2, 4, 8, 16}) and also evaluated at full depth (25 layers).

Optimization settings were selected before the main experiments by tuning the classifier on AP2-I and using those parameters for all other models. Optuna was used to search the parameter space to maximize the validation AUPRC. The hyperparameter search covered learning rates, weight decay, and the learning rate schedule, as well as the optimizer and warmup steps. On the basis of these results, we fixed weight decay at 0.01, retained AdamW, and used 1,000 warmup steps. Models were evaluated every 1,000 steps and the checkpoint with the best validation AUPRC was retained. Class imbalance between bound and unbound sequences was addressed by a class-balanced sampler: training examples were drawn with replacement in inverse proportion to their class frequency, yielding approximately balanced batches. The pretrained model core was adapted with the LPFT schedule (see **Downstream task fine-tuning strategies**) with the classification layers trained with a learning rate of 1e-5 and the model core learning rate was set to 1e-6 once the weights were unfrozen. Models without pretraining were trained from random initialization with a learning rate of 1e-6. For each of the three random seeds (11, 22, 33), the entire model was randomly initialized under that seed and, for the pretrained runs, only the shared model core weights were used as initializations. Consequently, the model without pretraining and the pretrained model’s classification layers began from identical weights for a given seed, and the two conditions differed only in the model core’s initialization (random vs. pretrained).

### Downstream task fine-tuning strategies

We evaluated four strategies for adapting the pretrained model core during classification. All three share the same learning rate (1e-5) for the classification layers, the same model core learning rate (1e-6), and the same number of training steps (20,000). “Linear probing then fine-tuning” (LPFT; used for the reported results) splits training into two phases. During the first phase, the convolutional model core is frozen and held in evaluation mode and only the classifier layers are trained. At the midpoint of training, the model core is unfrozen and the weights are allowed to be updated with a learning rate of 1e-6 (1/10 the learning rate of the classifier layers). In the “Full Frozen” fine-tuning method, the model core is frozen and kept in evaluation mode for the entire run and only the classification layers are trained. The “Lower Model Core Learning Rate” strategy does not freeze any layers, but the model core and classifier layers are assigned different learning rates. The model core trained with a learning rate of 1e-6, while the classifier layers trained with a learning rate of 1e-5. In the “None Frozen” setting, the model core shares the same learning rate as the classification head and is allowed to update normally for the full training run.

### Datasets

We used ChIP-seq datasets from multiple experimental sources and one ATAC-seq dataset (Toenhake et al., 2018) to train our classification models. We downloaded all publicly available ApiAP2 TF ChIP-seq data and a collection of histone modification and histone variant ChIP-seq data from the sources listed below. For each factor, MACS2 narrowPeak files (see above in **Reconstruction scores at *Pf*AP2-G2 motif-containing peaks**) were concatenated across available datasets, coordinate-sorted, and merged using bedtools merge to produce a single non-redundant peak set. Where a factor was referred to by multiple names in the literature (*Pf*AP2-P, also known as *Pf*AP2-11A), peak files from both sources were pooled prior to merging. We filtered out any dataset with fewer than 10 peaks on the test chromosome (chr 14). The remaining merged BED files were used as the positive set for binary classification. The ATAC-seq dataset and the chromatin states were processed as in Brown et al., (2026). The majority of the TF datasets came from one study (Shang et al., 2022): *Pf*AP2-L, *Pf*AP2-G4, *Pf*AP2-HS, *Pf*AP2-G3, *Pf*AP2-G5, PF3D7_0613800, *Pf*AP2-O2, PF3D7_1239200, PF3D7_1115500, and *Pf*AP2-O5. Some datasets were reported by both Shang et al. (2022) and other studies: *Pf*AP2-HC (Carrington et al., 2021; Shang et al., 2022), *Pf*AP2-G2 (Shang et al., 2022; S. Singh et al., 2021), *Pf*AP2-P (Shang et al., 2022; Subudhi et al., 2023), *Pf*AP2-EXP (Russell et al., 2022; Shang et al., 2022) and *Pf*AP2-LT (Bonnell et al., 2024; Shang et al., 2022). Two ApiAP2 TF datasets were sequenced by just one study: *Pf*AP2-I (Santos et al., 2017) and *Pf*AP2-G (Josling et al., 2020). *Pf*HP1 (Carrington et al., 2021; Shang et al., 2022) came from two studies and MORC from one (M. Singh et al., 2024). Downloaded histone mark data included H2A.Z, H3K4me3, and H3K9ac (Bartfai et al., 2010), H3K4me1 (J. Tang et al., 2020), and H3K18me1 (Reers et al., 2023). All chromatin states were reported in (Brown et al., 2026).

### Feature attribution with TPCAV

TPCAV v0.10.3 (https://github.com/seqcode/TPCAV) was used to train linear classifiers for the evaluated concepts. Position weight matrices (PWMs) for the DNA motif concepts were obtained from (Campbell et al., 2010) and MEME motif scans on ChIP-seq datasets. Motif concepts were evaluated by inserting 3, 6, or 12 motif instances into randomly sampled 512 bp genomic sequences. For TPCAV’s PCA transformation step, 120 examples were sampled from each concept and pooled across all concepts. Subsequently, 5,000 examples per concept were used to train the linear classifiers.

### Comparing to the MalariaSED-like model

We reimplemented the convolutional–recurrent architecture of MalariaSED (Wang et al., 2023) and compared it with our pretrained GPN classifiers across sixteen DNA-binding factors (**Supplemental Fig. 4**). Input sequences were one-hot encoded over A, C, G, T, and N and passed through two 1D convolutional layers of 512 filters each, followed by a nonlinear activation (ReLU or tanh, selected per factor during hyperparameter search), max pooling and dropout; a single bidirectional long short-term memory (LSTM) layer; and two fully connected layers producing a single logit. Both convolutional layers used 512 filters with a 9 bp kernel, matching the GPN encoder. Both pipelines used the merged ChIP-seq peak sets described above and the *P. falciparum* 3D7 reference assembly. We used the same validation and test chromosomes for both (chr 13 for validation and chr 14 for test).

Each architecture was trained on the example set appropriate to its published design. For the MalariaSED-like models, the genome was tiled into 200 bp bins following Wang, et al. (2023). A bin was labeled positive if at least 60% of its length overlapped a merged peak and negative if less than 20% of its length overlapped; bins with intermediate overlap were discarded. For each retained bin, we extracted a 1,000 bp sequence window centered on that bin, and the training set was augmented with the reverse complement of every window. Examples were partitioned into training, validation and test sets by chromosome as above. The GPN classifiers were trained as described above. The peak counts annotated in **Supplemental Fig. 4** are the numbers of merged ChIP-seq peaks falling on the training and test chromosomes and are therefore common to both architectures. The MalariaSED-like models were trained with three random seeds each.

Performance was quantified as AUPRC. For each factor, we report the mean and standard deviation of AUPRC across replicate seeds. For the cross-architecture comparison, MalariaSED-like models trained from scratch were compared with pretrained GPN LPFT models at full 25-layer depth. Differences were assessed with Welch’s two-sided two-sample t-test, which assumes neither equal variances nor equal replicate counts, applied to every factor. p-values were corrected across factors using the Benjamini-Hochberg procedure and differences were called significant at q < 0.05.

## Acknowledgements

Research reported in this publication was supported by NIGMS of the National Institutes of Health under award number R35GM144135 (to S.M.). Over 80% of the work described in this study was supported by this NIH award. A.S.B. acknowledges training opportunities (unfunded during this work) from the Eukaryotic Gene Regulation training grant (T32GM152354). A.S.B. was also supported by a GlaxoSmithKline Graduate Fellowship during part of this work. The content is solely the responsibility of the authors and does not necessarily represent the official views of the National Institutes of Health. The authors thank members of the Center for Eukaryotic Gene Regulation at Penn State for feedback on the work.

