## Supplemental Information for "An apicomplexan-specific foundation model improves prediction of regulatory genomic signals in *Plasmodium falciparum*"

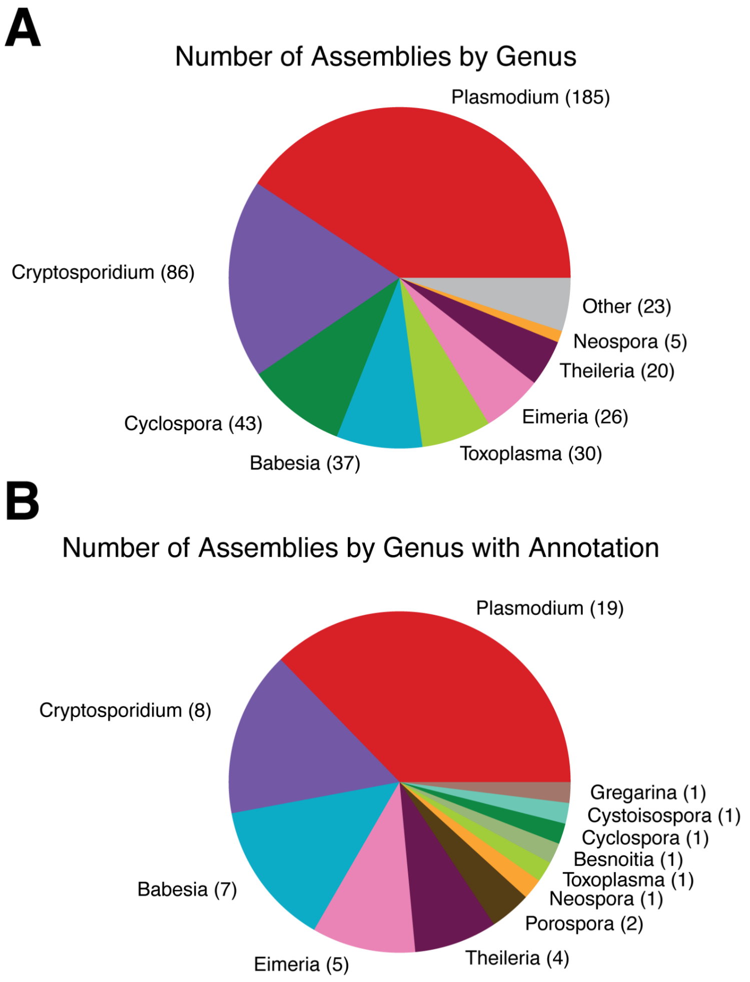


**Supplemental Figure 1.** Pie plots showing assemblies by genus. **A)** The number of each genus in the 455 assemblies used for the annotation-independent reconstruction models. Thirteen genera were merged into the "Other" category (23 assemblies total): Porospora: 4, Hammondia: 3, Nephromyces: 2, Cystoisospora: 2, Gregarina: 2, Sarcocystis: 2, Besnoitia: 2, Ascogregarina: 1, Cytauxzoon: 1, Cardiosporidium: 1, Ophryocystis: 1, Hepatocystis: 1, and Haemoproteus: 1. **B)** The number of each genus in assemblies with gene annotations.

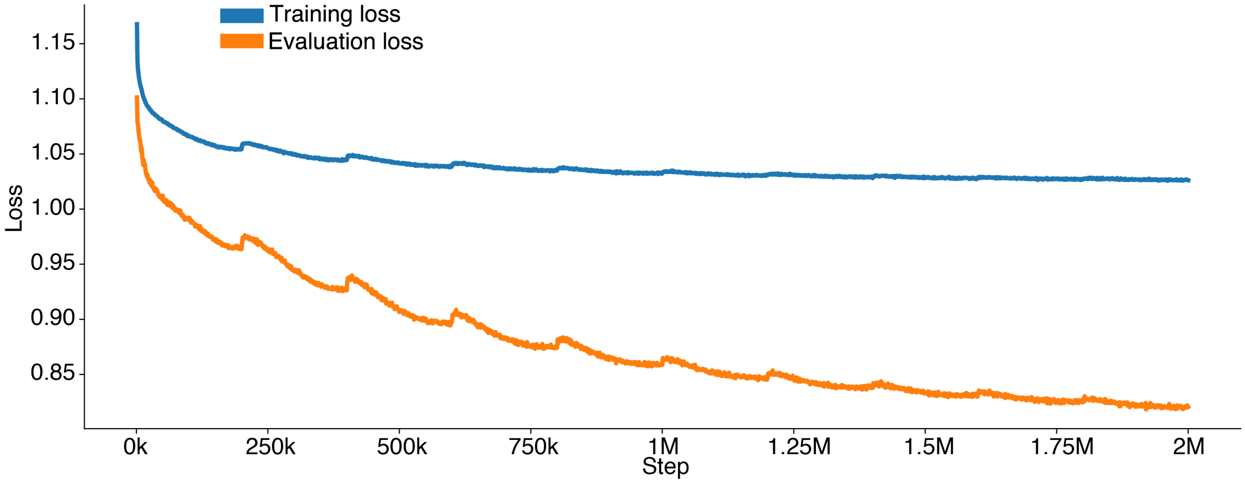


**Supplemental Figure 2.** Training and evaluation loss for the "Whole Genome 455A 2M steps" model.

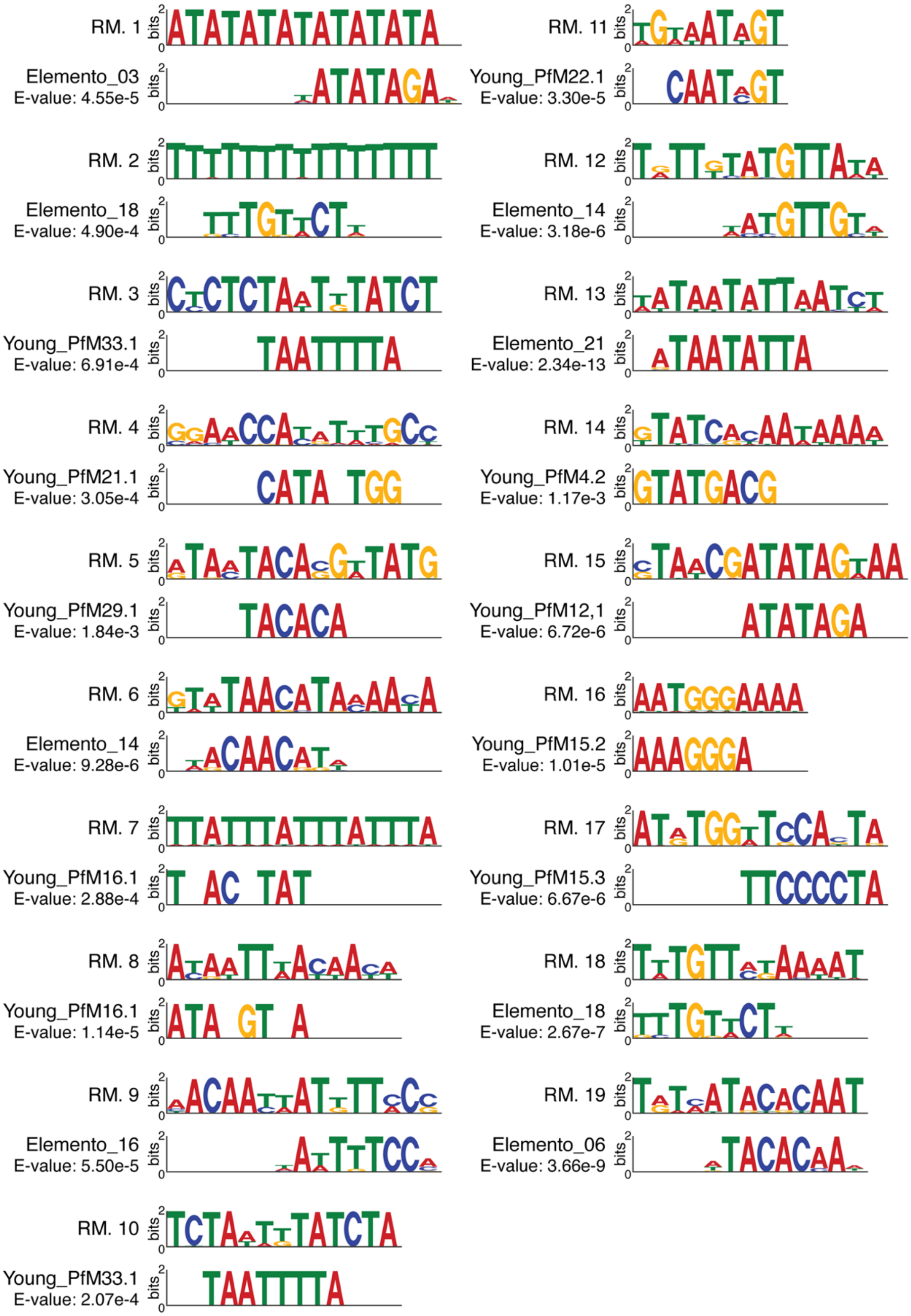


**Supplemental Figure 3.** *De novo* motifs discovered in highly reconstructible promoter regions and matches to predicted motifs (Elemento et al., 2007; Young et al., 2008).

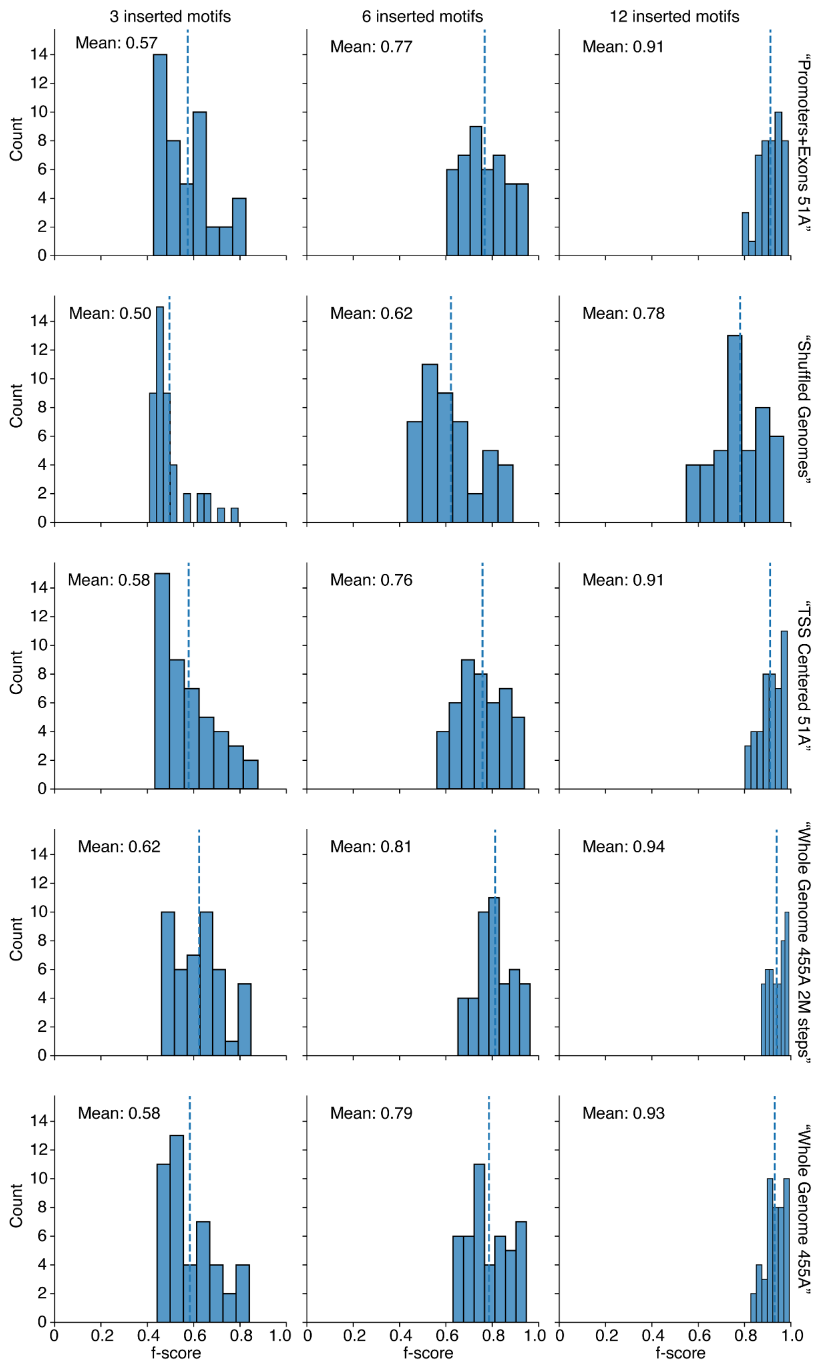


**Supplemental Figure 4**. TPCAV classifier performance distributions for motif concepts in each model. Plots show F-score distributions of motif concepts under different numbers of motif insertions in concept example sequences.

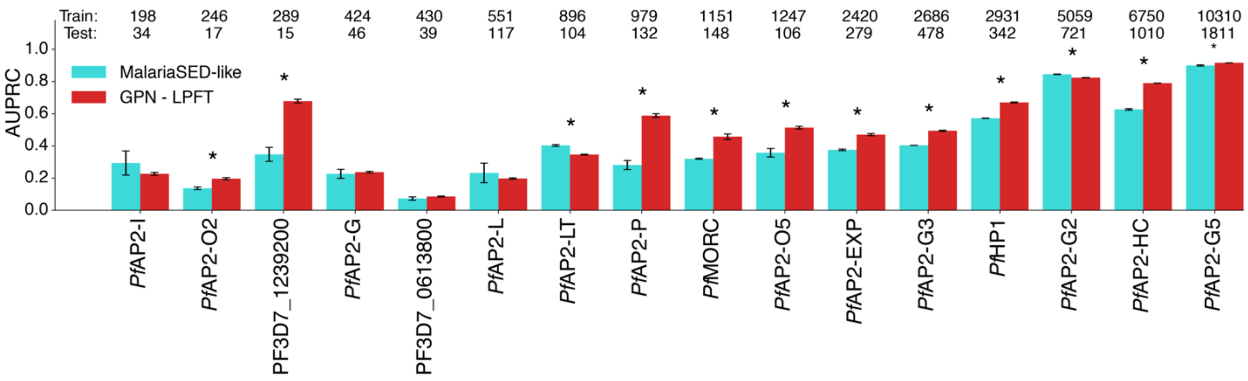


**Supplemental Figure 5**. Cross-architecture comparison of AUPRC on the held-out test chromosome (chr 14) for sixteen *P. falciparum* DNA-binding factors. Cyan bars, MalariaSED-like classifiers trained from randomly initialized weights; red bars, GPN classifiers using the full 25-layer "Whole Genome 455A 2M steps" model core with LPFT fine-tuning. Factors are ordered left to right by increasing numbers of ChIP-seq peaks on the training chromosomes; the numbers of peaks on the training and test chromosomes are shown above each pair of bars (training on top, test below). Bar heights are the mean of three random seeds and error bars show ± 1 standard deviation. Asterisks mark factors at which the two architectures differ significantly (Welch's two-sided two-sample t-test on replicate AUPRC values, Benjamini–Hochberg corrected across factors, $q < 0.05$). The pretrained GPN model is higher at every marked factor except AP2-LT and AP2-G2, where the MalariaSED-like model is higher.
